# Monitoring regional astrocyte diversity by cell-type specific proteomic labeling *in vivo*

**DOI:** 10.1101/2022.06.07.495069

**Authors:** Priyadharshini Prabhakar, Rainer Pielot, Peter Landgraf, Josef Wissing, Anne Bayrhammer, Marco van Ham, Eckart D. Gundelfinger, Lothar Jänsch, Daniela C. Dieterich, Anke Müller

## Abstract

Astrocytes exhibit regional heterogeneity in morphology, function and molecular composition to support and modulate neuronal function and signaling in a region-specific manner. To characterize regional heterogeneity of astrocytic proteomes of different brain regions we established an Aldh1l1- MetRS^L274G^ mouse line that allows cell-type specific labeling of newly synthesized proteins *in vivo* and analyzed astrocytic proteins from four different brain regions by mass spectrometry. Identified proteins are specific for astrocytes and show a high overlap with proteins compiled in ‘AstroProt’, a newly established database for astrocytic proteins. Gene enrichment analysis reveals high overlap among brain regions, and only subtle changes in abundances of key astrocytic proteins for hippocampus, cortex and striatum. However, the cerebellar proteome stands out with proteins being associated with calcium signaling or bipolar disorder. Subregional differences of translation dynamics in single hippocampal astrocytes indicates distinct subregional heterogeneity and highlights the applicability of our toolbox to study dynamic astrocytic proteomes *in vivo*.

## Introduction

Astrocytes are the most abundant glia cell type in the mammalian brain and as they shape and modulate neuronal signaling it is essential to understand their multifaceted roles in order to unravel the complexity of neuronal signaling processes. Astrocytes provide metabolic support to neurons, they shape and synchronize neuronal activity and they form contacts with neurons by fine astrocytic processes to form the so-called multipartite synapse (Durkee & Araque, 2019; Verkhratsky & Nedergaard, 2018; Witcher et al., 2007). At the multipartite synapse, astrocytes support, sense and modulate synaptic activity and thereby contribute to synaptic signaling (Durkee & Araque, 2019; Verkhratsky & Nedergaard, 2018). This impact is not only restricted to local effects but is far-reaching as astrocytes shape neuronal circuit activity and network oscillations (Buskila et al., 2019; Mederos et al., 2018).

With their fundamental role in multifaceted brain processes it is no surprise that astrocytes influence the onset and progression of neuronal disorders ranging from neurodegenerative diseases such as Alzheimer’s disease to mood disorders (Phatnani & Maniatis, 2015; B. Zhou et al., 2019; X. Zhou et al., 2019). Again, the impact by astrocytes is context-dependent and influences the tight interrelationship with neurons. For example, a loss of synaptic support or modulation of synaptic activity by astrocytes was recognized to contribute to neurodegenerative diseases (Blanco-Suárez et al., 2017).

Even though, astrocytes are undoubtedly influential in many ways, little is known about regional specialization of astrocytic support and modulative capacities. Astrocytic subtypes with regional specialization could, for example, shape selected functions in network activity or synaptic signaling. Indeed, the notion that astrocytes show intra- and interregional heterogeneity in morphology, molecular composition and physiology has been substantiated manifold (B. E. Clarke et al., 2021; Khakh & Deneen, 2019; Pestana et al., 2020). Specialized astrocytic subtypes were identified in certain brain regions, such as Müller cells in the retina or Bergmann glia cells in the cerebellum supporting the idea that astrocytes comprise a group of cells that have confined function in brain physiology (Verkhratsky & Nedergaard, 2018). Further identification of regional astrocytic subtypes had been also correlated to morphological and physiological differences. Astrocytes from distinct brain regions revealed differences in Ca^2+^ signaling, electrophysiological properties or modulation of synapse formation, for example (Chai et al., 2017; Lin et al., 2017; Morel et al., 2017). This physiological complexity reaches further into subregions of the brain. Cortical or striatal subregional astrocytes, for instance, display differential Ca^2+^ signaling in response to neuronal activity or layer-specific morphological characteristics (Farhy-Tselnicker et al., 2021; Lanjakornsiripan et al., 2018). Although transcriptomic studies provide the sensitivity needed to unravel the molecular composition of cells in restricted areas and allow even single-cell analysis (Batiuk et al., 2020; Bayraktar et al., 2020; Doyle et al., 2008; Zeisel et al., 2015) they might not fully reflect the cellular proteome and further analysis of proteome dynamics in astrocytes complements the characterization of astrocytic complexity and substantiates a clear categorization and characterization of regional astrocyte subtypes. First studies indeed aimed to assemble a complete picture of the molecular composition of regional-diversified astrocytes by a combination of transcriptomic and proteomic techniques (Chai et al., 2017; Sharma et al., 2015).

To complement the toolbox for astrocytic proteome analysis by an *in vivo* metabolic labeling approach, we generated a transgenic mouse line containing a floxed-STOP version of a GFP-2A-MetRS^L247G^ construct for conditional expression that is crossed with the tamoxifen-inducible Cre-driver line Aldh1l1-CreERT2 to metabolically label the astrocyte-specific proteome *in vivo* (Alvarez-Castelao et al., 2017; Winchenbach et al., 2016). Subsequent activation of the Cre-recombinase by tamoxifen leads to the astrocyte-specific expression of the mouse methionyl-tRNA synthetase version MetRS^L274G^ that includes an expanded methionine binding site to enable the activation and subsequent incorporation of the methionine surrogate azidonorleucine (ANL) into nascent peptides during translation (Erdmann et al., 2015; Müller et al., 2015). ANL endows proteins with a new functionality that can be used in downstream applications for click chemistry-based approaches like Bioorthogonal non-canonical amino acid tagging (BONCAT) or Fluorescent non-canonical amino acid tagging (FUNCAT) (Dieterich et al., 2006, 2010). Hereby, either a biotin- or fluorescence-bearing tag is coupled to the Azide-group in the side chain of ANL by ‘click reaction’ carried out on either protein lysates or brain tissue slices, respectively.

Herein, we confirm the specificity of this method for *in vivo* labeling and analysis of newly synthesized proteins with astrocytic origin in four areas of the mouse brain, as supported by the high overlap with AstroProt entries and the identification of several key astrocytic proteins. Cerebellar astrocytes, as compared to astrocytes from the other three regions studied, are unique in displaying a clear abundance of calcium signaling pathway components and a significant correlation with genes associated with bipolar disorder. Our work provides a unique tool to label astrocytic proteins *in vivo* and to disclose dynamic astrocytic proteome changes in physiology, behavior and disease.

## Results

### Astrocyte-specific expression of MetRS^L274G^ allows labeling of newly synthesized proteins *in vivo*

The tight interconnection of neurons and astrocytes in the brain with astrocytes forming a variety of cell contacts with neuronal neurites and synapses makes it technically challenging to study the astrocytic proteome, as this requires disentangling astrocytes from neuronal cell populations. To be able to separate astrocytic proteins from the neuronal proteome, we used cell type-selective metabolic labeling to tag, isolate and analyze proteins synthesized by astrocytes *in vivo*. To this end, we crossed homozygous R26-GFP-2A-MetRS^L274G^ mice (further referred to as MetRS^L274G^ mice) with the inducible Cre-driver line Aldh1l1-CreERT2 allowing the conditional expression of the mouse Methionyl-tRNA synthetase^L274G^ (MetRS^L274G^) and GFP exclusively in Aldh1l1-expressing astrocytes **(Figure 1A)** (Alvarez-Castelao et al., 2017; Winchenbach et al., 2016). Here, enlargement of the MetRS methionine binding pocket enables azidonorleucine (ANL) coupling to ^Met^tRNA, random incorporation of ANL into nascent proteins during translation and the subsequent application of FUNCAT or BONCAT for visualization or isolation of tagged proteins **(Figure 1B)** (Dieterich et al., 2006, 2010).

**Figure 1.**
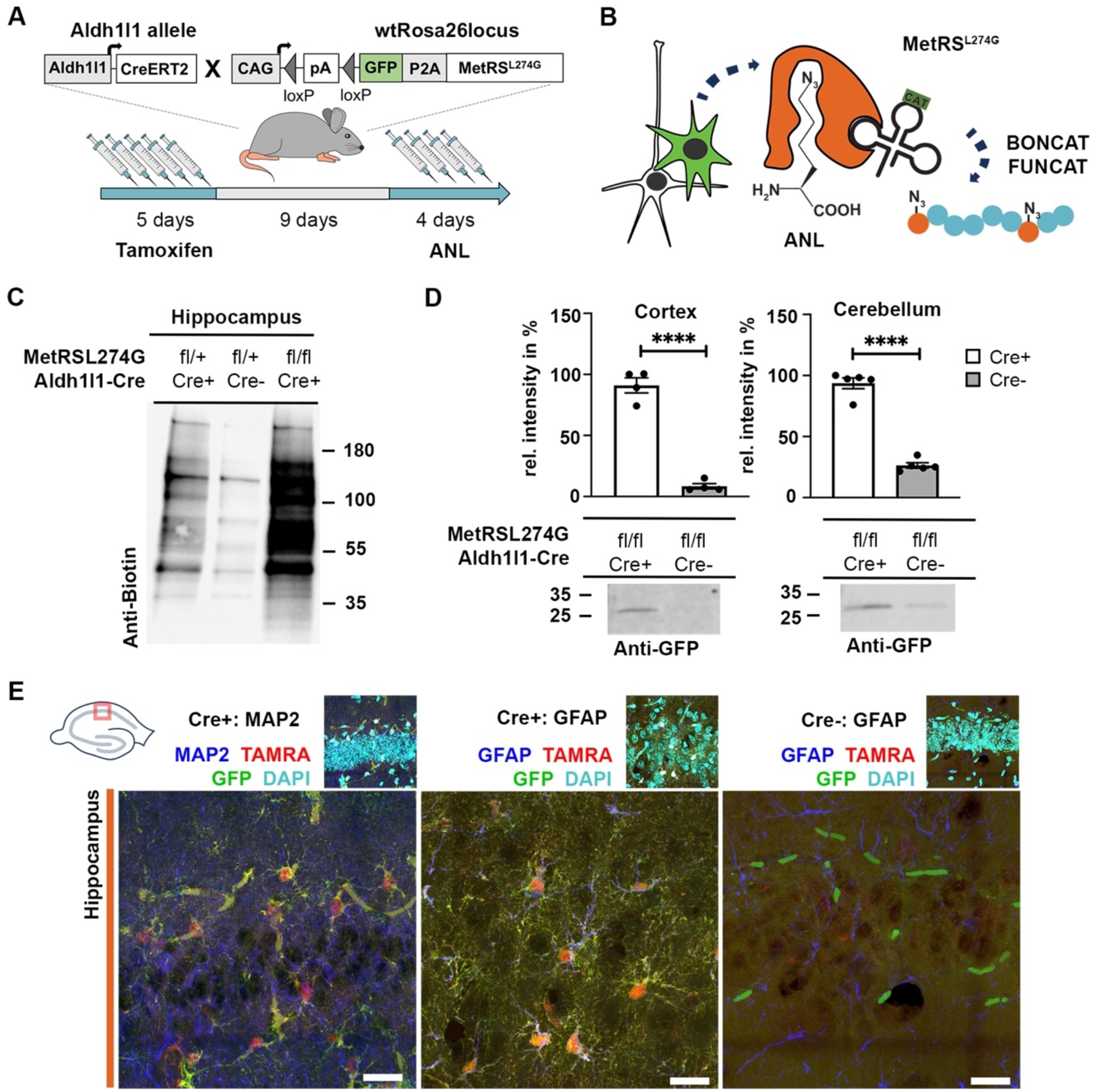
Astrocyte-specific labeling of newly synthesized proteins *in vivo*. **(A)** Breeding scheme of mice driving CreERT2 from the astrocyte-specific Aldh1l1 promoter with mice carrying the inducible transgene for expression of GFP and MetRS^L274G^. Protein labeling in astrocytes is achieved by the indicated schedule of tamoxifen followed by azidonorleucine (ANL) injection (see also Supplemental Figure 1) **(B)** MetRS^L274G^ expressed in astrocytes enables coupling of ANL to Met-tRNA, integration into nascent proteins and tagging by ‘click reaction’. **(C)** Immunoblotting of tissue derived from mice homozygous (MetRS^L274G; fl/fl^) or heterozygous (MetRS^L274G; fl/+^) with or without Cre expression (Cre+/-) after chemical ‘click reaction’ with a biotin tag. **(D)** GFP reporter expression was quantified using immunoblotting from cortical or cerebellar tissue samples of Cre+ or Cre- MetRS^L274Gfl/fl^ mice. (mean ± SEM; n=4-5/group, technical replicate (tr)=2; two-tailed t-test; ****: p<0.0001). **(E)** Hippocampal slices were tagged with a fluorescence tag (TAMRA) by ‘click reaction’ and processed by immunohistochemistry (scale bar=20µm). Neither GFP nor TAMRA signal was evident in Cre- derived slices (autofluorescent erythrocytes generate unspecific signal in the GFP channel).

Optimal labeling efficiency was achieved after a daily i.p. injection of tamoxifen for five days and, after a 9-days break, by i.p. injection of ANL for four consecutive days **(Figure 1A)**. Monitoring of health status and weight of mice being able to integrate ANL into proteins did not reveal any noticeable abnormalities **(Supplemental Figure 1)**. We further controlled for specificity of ANL labeling applying BONCAT, where brain tissue lysates of mice that are homozygous for the MetRS^L274G^ transgene (MetRS^L274G; fl/fl^) produce distinct biotin signals after ‘click reaction’ with a biotin-tag. In contrast, samples derived from MetRS^L274G^ mice lacking Cre expression (Cre-) were almost devoid of biotin signal in immunoblotting **(Figure 1C).** As MetRS^L274G; fl/fl^ generate stronger labeling compared to heterozygous MetRS^L274G; fl/+^, only MetRS^L274G; fl/fl^ were used hereinafter, further referred to as MetRS^L274G^. Quantification of GFP reporter expression as exemplified for cortical and cerebellar tissue proves specificity of induced MetRS^L274G^ expression by tamoxifen **(Figure 1D).**

We also controlled for cell-type specificity of astrocytic labeling using FUNCAT in brain slices, where a fluorescent TAMRA-alkyne tag after ‘click’ reaction’ visualizes ANL-bearing proteins **(Figure 1E)**. We found clear co-labeling of the GFP reporter and the astrocyte marker GFAP in Cre+ MetRS^L274G^ mice, but not with the neuronal marker MAP2, proving GFP expression is specific for astrocytes. Also here, the TAMRA-signal was exclusively found in GFP-positive cells of Cre+ MetRS^L274G^ mice confirming cell-type specificity of *in vivo* protein labeling in astrocytes via MetRS^L274G^ and ANL.

### Analysis of astrocytic proteomes in subregions of the mouse brain

To unravel potential regional differences in astrocytic proteomes, we performed *in vivo* labeling of astrocytic proteins in 8-12-week-old mice. Three Cre+ MetRS^L274G^ and two Cre-MetRS^L274G^ mice were sacrificed 24 hr after the last ANL injection and brains were dissected into cortex (Cx), hippocampus (HC), striatum (Str) and cerebellum (Ceb). Protein extracts were subjected to ‘click reaction’ either with a biotin-tag for immunoblot analysis or coupled onto agarose alkyne beads for subsequent isolation by affinity purification and identification by liquid chromatography–mass spectrometry (LC-MS/MS) analysis and label-free quantification **(Figure 2A)**.

**Figure 2.**
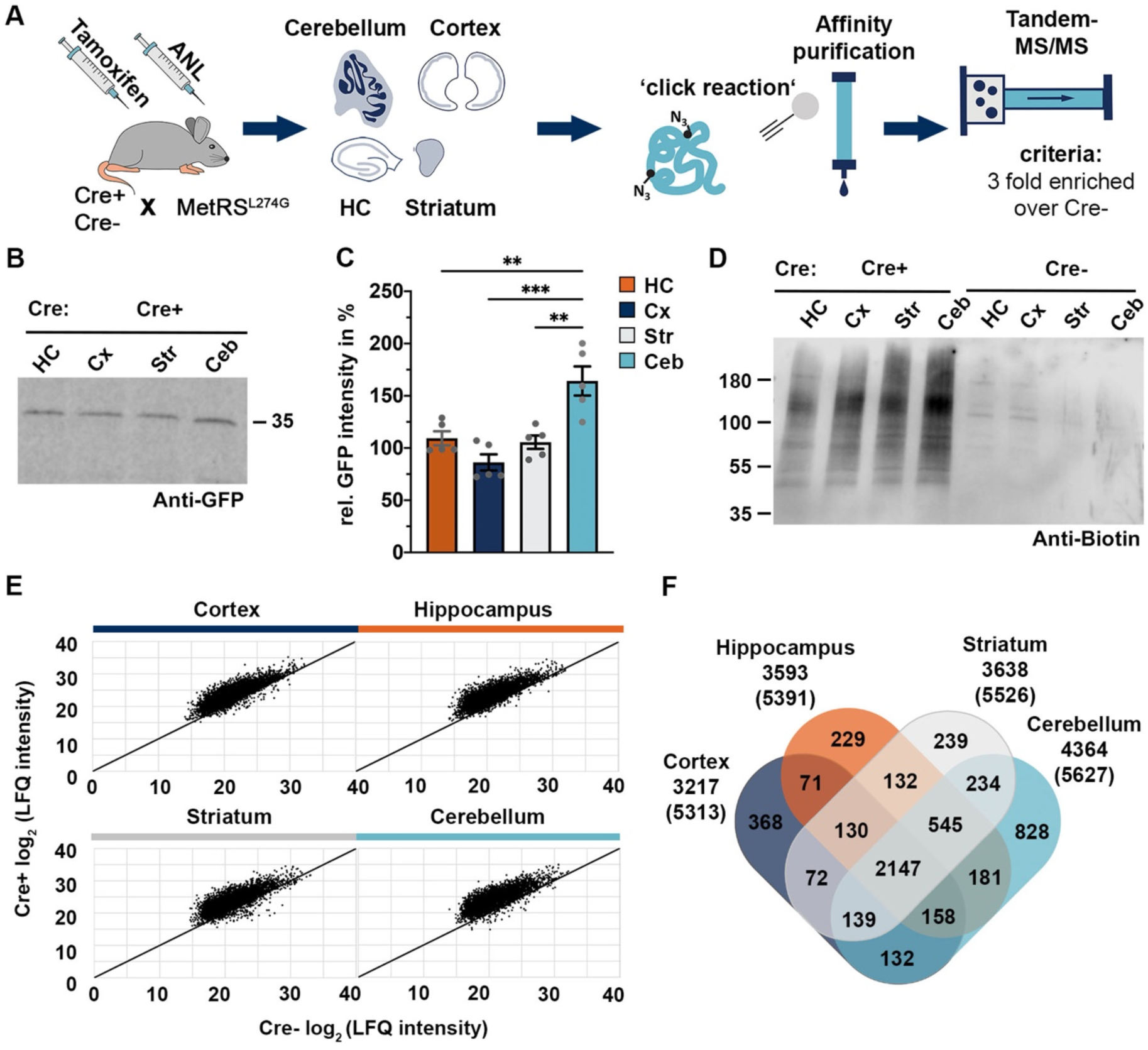
Regional analysis of the newly synthesized astrocytic proteome. **(A)** Steps to obtain regional proteomes. Tissue lysates from three Cre+ or two Cre- MetRS^L274G^ mice were ‘clicked’ directly onto alkyne resins, followed by a trypsin peptidase digest to release peptides from beads for LC/MS-MS analysis. **(B)** GFP expression in brain regions compared by immunoblotting and quantification **(C)** shows similar levels in hippocampus (HC), cortex (Cx), striatum (Str) but enhanced expression in the cerebellum (Ceb) (mean ± SEM; n=5/group, tr=2; One-Way ANOVA; multiple comparisons; **: p<0.01; ***p<0.001; F(3)=13.25, p=0.0001). **(D)** Brain tissue of either Cre+ or Cre- MetRS^L274G^ mice were tagged with a biotin-alkyne tag and visualized by immunoblotting (See also Figure S2). **(E)** Scatter plot of MS-MS data of each brain region reveals higher protein log2 label-free quantitation (LFQ) intensities for Cre+ samples compared to Cre- samples for all four brain regions analyzed. **(F)** Venn diagram shows numbers of ≥3-fold enriched proteins for each brain region. (numbers in brackets refer to total number of identified proteins per region. See also Supplemental Figure 2 and 3).

To control for comparability of samples derived from different brain regions, we quantified GFP reporter expression of Cre+ animals by immunoblotting **(Figure 2B, C).** Whereas GFP levels are similar in the hippocampus, cortex and striatum, we found a significantly higher GFP expression in the cerebellum. We further analyzed the MetRS^L274G^ dependent protein synthesis capacities in selected regions utilizing the alkyne-bearing biotin tag in ‘click reaction’ and detected robust biotin signals in all protein extracts from MetRS^L274G^ mice **(Figure 2D).** To evaluate comparability of samples, we performed dot-blot assays with biotinylated protein extracts and found similar levels of biotin-tagged proteins in all four brain regions tested **(Supplemental Figure 2B).** We further controlled ANL integration by immunohistochemistry in all four brain regions applying a TAMRA- tag **(Supplemental Figure 2A)**.

Purified ANL-tagged protein samples from all four brain areas were run in triplicates in LC-MS/MS and label free quantification (LFQ) was used to compare intensities of single proteins. Identified brain region-specific protein sets were characterized by plotting the log2 of label-free quantitation intensities of Cre+ MetRS^L274G^ samples with their respective Cre- control. We found clearly enhanced log2 intensities of Cre+ derived proteins in scatter plots **(Figure 2E),** proving the enrichment for Cre+ MetRS^L274G^ samples in affinity purification for all four brain regions. Specificity is further supported by scatter plots that proof low sample variation among biological replicates within each group **(Supplemental Figure 3).** We further observed similar protein classes when compared to proteins identified by Sharma et al.(Sharma et al., 2015) **(Supplemental Figure 2C)**. As valid identification criterion, we defined proteins to have at least a 3-fold enriched abundance in Cre+ MetRS^L274G^ mice over Cre- control samples and compared proteins from all four datasets in a Venn diagram **(Figure 2F)**. We identified 3,217 proteins in the cortex (60.5% of the complete list), 3,593 in the hippocampus (66.6%), 3,638 in the striatum (65.8%) and 4,364 proteins in the cerebellum (77.6%) to meet this criterion. A high overlap of about 60% of identified proteins is present in all four brain regions indicating that the overall sets of proteins do not vary grossly among brain regions. Also, the number of proteins that are unique for one brain region are rather small, we find 368 unique proteins in the cortex, 229 in the hippocampus, 239 in the striatum and 828 in the cerebellum.

### Assessment of regional astrocytic proteomes utilizing the AstroProt database

To furthermore validate the origin of the identified proteins from various brain regions, we compared our datasets with data from a previous study (Sharma et al., 2015). Calculation of the percentage of astroglial and neuronal marker (677 or 713 proteins) that were extracted from Sharma *et al*. within our datasets shows an enrichment of astroglial proteins in all four brain-specific proteomes **(Figure 3A)**. The relative proportion of astroglial marker ranges from 31,7% (cortex) to 39,6% (cerebellum), whereas the relative proportion of neuronal marker ranges only from 8,4% (cortex) to 18,5% (cerebellum).

**Figure 3.**
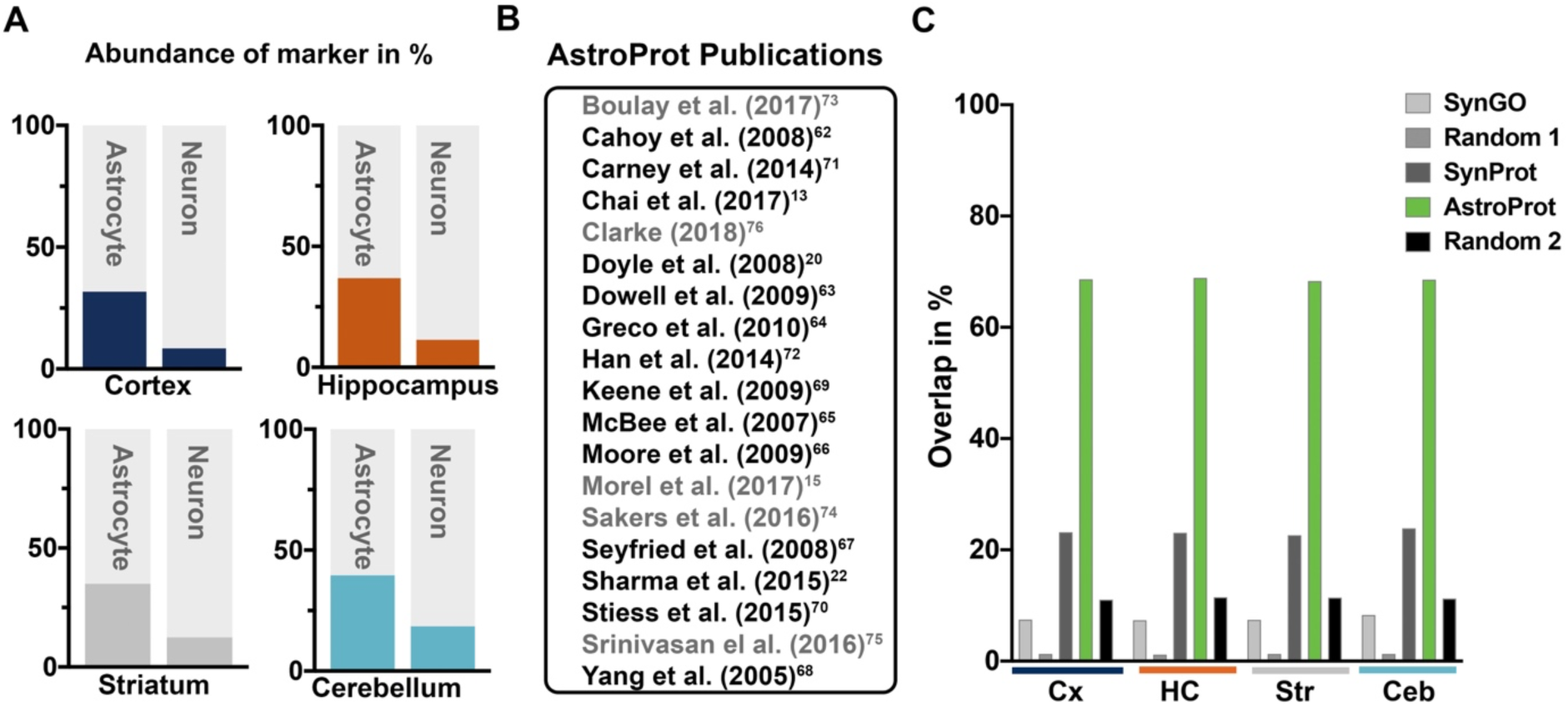
Assessment of regional astrocytic proteomes utilizing AstroProt. **(A)** Relative abundances of previously defined cell-type specific protein marker in area-specific proteomes. Depicted are fractions of marker found relative to the total number of either neuronal (677 proteins) (Sharma et al., 2015) or astrocytic marker (713 proteins) (Sharma et al., 2015) used in this analysis. Astroglial marker are enriched in all four brain areas. **(B)** List of publications integrated into AstroProt database. Publications in grey refer to studies identifying astrocytic ribosome-bound mRNA. **(C)** Identified datasets for each brain region were compared with AstroProt (mouse protein entries only), the synapse-specific databases SynGO and SynProt and two random lists (Random 1: 1,044 proteins; Random 2: 9,659 proteins). The relative overlap is shown between the area-specific proteomes (only 3-fold enriched) and these databases. (Cx: cortex, HC: hippocampus, Ceb: cerebellum, Str: striatum. See also Supplemental Figure 4).

To further confirm the specificity of astrocytic datasets, we generated a database compiling 13,665 protein entries from studies characterizing the mouse, human and rat astrocytic proteome and selected ribosome-bound mRNA, which we designate AstroProt. To establish AstroProt, information about astroglial proteins and ribosome-bound mRNAs were extracted from studies as listed in **Figure 3B** and provided as an additional database in the SynProt portal (https://www.synprot.de) (Pielot et al., 2012).

We compared the four datasets with 9,705 mouse proteins from AstroProt (excluding translation-active mRNA studies and proteomics studies from rat) and 2,316 mouse proteins from the database SynProt (Pielot et al., 2012). This database includes primarily neuronal proteins identified in synaptic junctional proteomes. Indeed, all four datasets overlap to very similar extent with AstroProt mouse proteins (Cx: 68.6%; HC: 68.9%; Ceb: 68.8%; Str: 68.3%), whereas the overlap with SynProt ranges from 22,6% (Str) to 23,9% (Ceb) **(Figure 3C)**. In addition, we compared datasets with synaptic mouse proteins from the expert-curated gene ontology Project SynGO (www.syngoportal.org) (Koopmans et al., 2019) generating an overlap of less than 10% identical entries. To exclude statistical effects, we compared our datasets with random protein lists of the mouse proteome with different sizes that were compiled by an inhouse script from the mouse proteome at uniport.org resulting in minor overlap of 1,2% (Ceb)-1.3% (Cx) (Random1) or 11.0% (Cx)-11.5% (HC) (Random2) **(Figure 3C)**. Due to a higher number of proteins in “Random2”, the possibility of overlap is statistically higher. This reconfirms that all four datasets are primarily of astrocytic origin.

AstroProt is accessible after registration at SynProt portal (www.synprot.de) (Pielot et al., 2012). This includes a web-based interface for an easy retrieval, where a single protein or a group of proteins can be searched e.g. for their name(s), parts of their sequence and further information, such as purification methods, age of cells and fractions (e.g. secretome or astrocytic protrusions) with optional selection of ribosome-bound mRNA **(Supplemental Figure 4)**.

### Region-specific gene ontology (GO) and KEGG enrichment analysis

To detect prominent biological processes and pathways in our datasets we performed gene enrichment analyses for all four brain regions. Analysis of ≥3-fold enriched proteins using inhouse scripts and GO of biological processes and subsequent plotting in a Venn diagram reveals a high similarity of brain regions with 44 biological processes that are commonly detected in all four brain regions **(Figure 4A; Supplemental Figure 5A)**. A detailed analysis of these biological processes revealed also uniquely enriched processes for all brain regions. Of those, we extracted the 10 GO annotation terms with the highest significance (Top 10) and identified processes concerning “Protein transport”, “Protein stabilization” and “Apoptosis” that are common in all regions or processes like “Cell migration” significantly enriched, for example, in the hippocampus (**Figure 4B).** For a better visualization of significantly enriched biological processes, we analyzed ≥3-fold enriched proteins with Cytoscape/ClueGO **(Figure 4C)**. The comparison of identified clusters shows that most of the enriched processes are common in all regions (e.g. processes concerning intracellular transport and cytoskeleton organization) or shared **(Supplemental Figure 5B)**. Again, some processes were found exclusively in one region like “mRNA metabolic processes” in the cortex. Interestingly, the functionality of the cerebellar proteome seems to differ from the other regions, as we identify several unique processes, such as “Neuron projection development” or “Cell junction organization” exclusively in this region.

**Figure 4.**
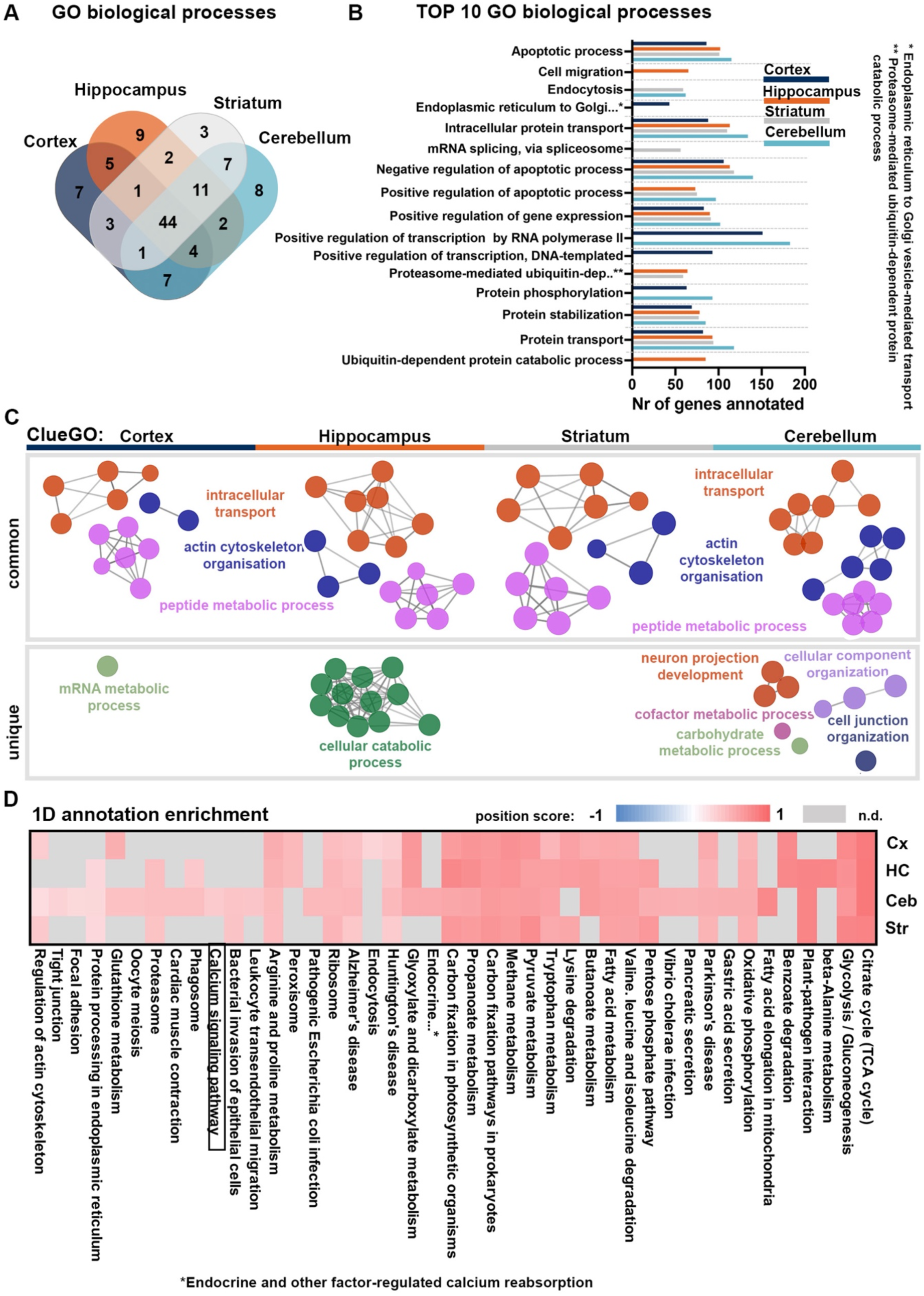
Region-specific gene ontology (GO) and KEGG enrichment analysis. **(A)** Significant GO terms of biological processes were determined using inhouse scripts and the number of significantly enriched GO terms were plotted for each analyzed brain region in a Venn diagram. The majority of significantly enriched GO terms is shared by more than one or even all brain regions. See also Figure S5 for lists of significant GO terms. **(B)** Top10 of significantly enriched GO terms concerning biological processes for each brain region. **(C)** Datasets of enriched proteins (≥3-fold change) for each region were analyzed by Cytoscape/ClueGO (GO biological process, only experimental evidence; p≤0.001; increasing node size denotes an enhanced significance). Shown are processes common for all brain regions or unique processes (for processes shared with two or three regions see also Supplemental Figure 5B). **(D)** Matrix of KEGG annotation terms enriched in different brain regions after clustering shows a prevalence in metabolic pathways with color-coded position scores after 1D annotation enrichment (cortex (Cx), hippocampus (HC), striatum (Str), cerebellum (Ceb). Gray fields represent lack of significant pathways.

For a quantitative gene enrichment analysis, we used the 1D enrichment function of Perseus (Cox & Mann, 2012). Here, the abundance of proteins generates a position score of a certain pathway, visualizing the enrichment. **Figure 4D** shows significantly enriched KEGG pathways for all four brain regions with metabolic pathways (e.g. “Citrate cycle”) being most prominent. Most of those pathways that were significantly enriched in only one brain region were found mainly in the cerebellum, indicating again a distinct functionality of cerebellar astrocytes.

### Astrocytic proteins enriched in regional proteomes

To further analyze region-specific abundances of proteins in different brain regions, we performed a cluster analysis using Perseus (Cox & Mann, 2012)**. Figure 5A** shows the first 5 clusters consisting of 47 proteins (complete heatmap in **Supplemental Figure 6A**). Among these proteins we find Glutamine synthetase (*Glul*), for example, that plays a pivotal role in the synaptic glutamate-glutamine cycle (Tani et al., 2014) or the enzyme Fructose-bisphosphate aldolase C (*Aldoc*) that, as part of the glycolytic pathway, contributes to neuronal energy supply (Deitmer et al., 2019). Whereas most enriched proteins were identified in all or in at least two brain regions with medium intensity variation, the dendrogram depicts again the special position of the cerebellum, as the cerebellar clusters are most different from other brain regions. We further compared brain regions pairwise, displaying significantly different protein expression abundances using volcano plots **(Supplemental Figure 6B).**

**Figure 5.**
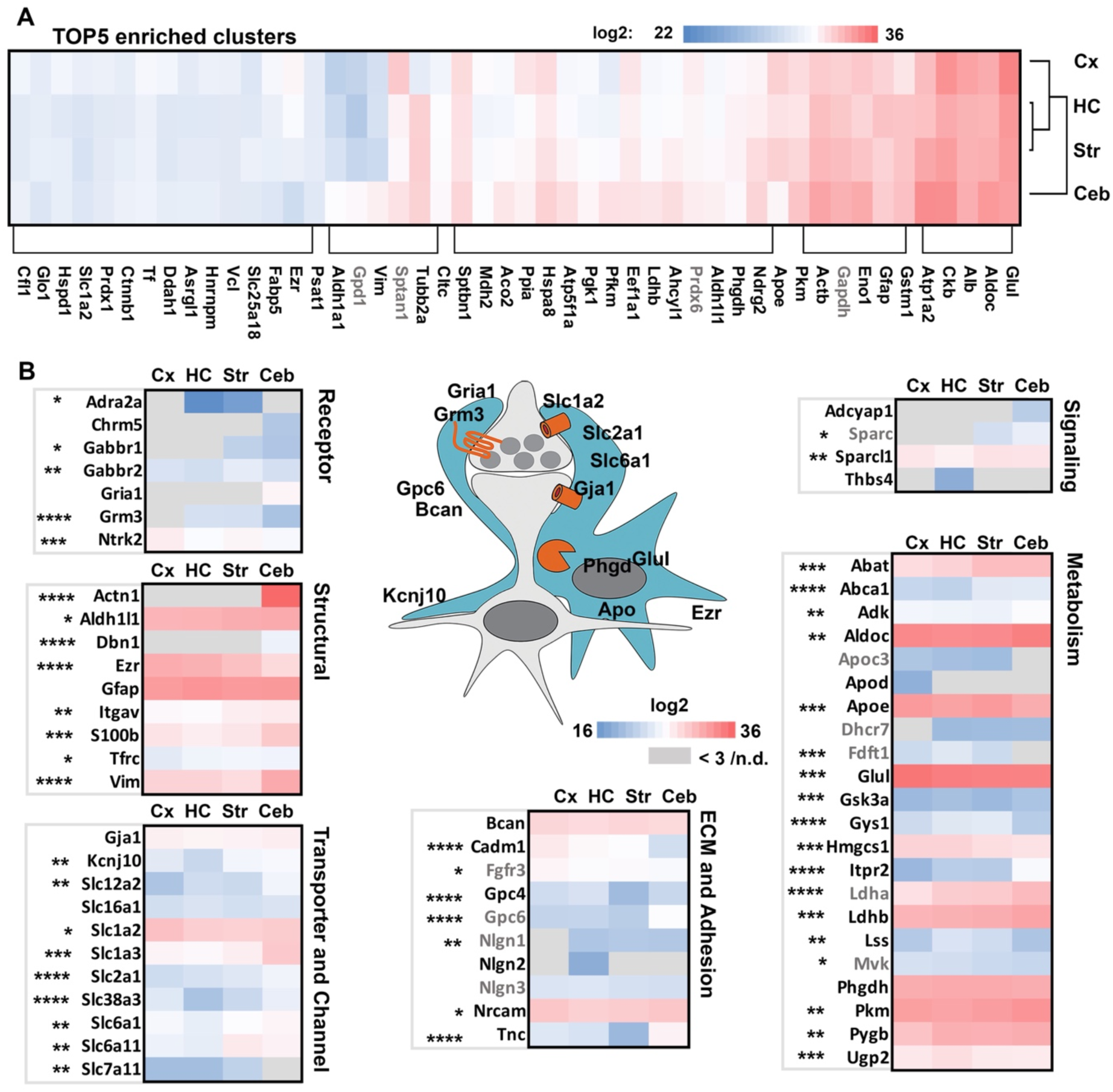
A variety of astrocytic proteins is enriched in regional proteomes. **(A)** Cluster analysis of protein intensities was performed with all four datasets and the top 5 enriched clusters are presented with color-coded log2 intensities. Gene names in black refer to reviewed protein entries in UniProt. **(B)** Prominent astrocytic proteins involved in modulation of synaptic activity or neuronal function were selected (as exemplified in the graph) and categorized into functional groups like receptors or transporters. The log2 intensities of 3-fold-enriched candidates were plotted for each brain region analyzed. Gene names in black refer to reviewed protein entries in UniProt. Gray fields represent lack of protein identification or less than 3-fold enrichment in this region. One-way ANOVA was performed to compare abundances in hippocampus (HC), striatum (Str), cortex (Cx) or cerebellum (Ceb) (*:p<0.05; **: p<0.01; ***p<0.001; ****p<0.0001). Gene names in black refer to reviewed protein entries in UniProt. (See also Supplemental Figure 6).

To further evaluate the robustness of data and identify differences in protein abundance in these four brain regions, we specifically selected proteins with a well characterized role in astrocytes focusing on their functional contribution to support and modulate neuronal physiology and signaling (Allen, 2014). Mean log2-intensities of 3-fold enriched proteins were displayed with color-coded intensities and statistically relevant differences of log2 intensities were compared among brain regions. Among these key astrocytic proteins were transporters for the uptake of neurotransmitters, such as Eaat2 (*Slc1a2*) **(Figure 5B).** Also, extracellular matrix proteins like Brevican (*Bcan*) as well as proteins with signaling function such as Sparc-like protein 1 (*Sparcl1*) were identified. Especially candidates of metabolic pathways were among those proteins with high log2 intensities and vastly represented in analyzed brain regions with subtle, but often significant differences in log2 intensities (**Figure 5B: “**Metabolism”).

We confirmed the expression of D-3-phosphoglycerate dehydrogenase (*PHGDH*) and Glycogen synthase 1 (*Gys1*), as representatives for metabolic enzymes, as well as Sparc-like protein1 (*Sparcl1*) by immunohistochemistry on brain slices derived from all four brain regions of Cre+ MetRS^L274G^ mice **(Figure 6)**. Indeed, immunoreactivity for those candidates was detected in astrocytes expressing GFP with expression levels resembling those predicted by proteomics data. Again, these results support the specificity of astrocytic origin of our datasets.

**Figure 6.**
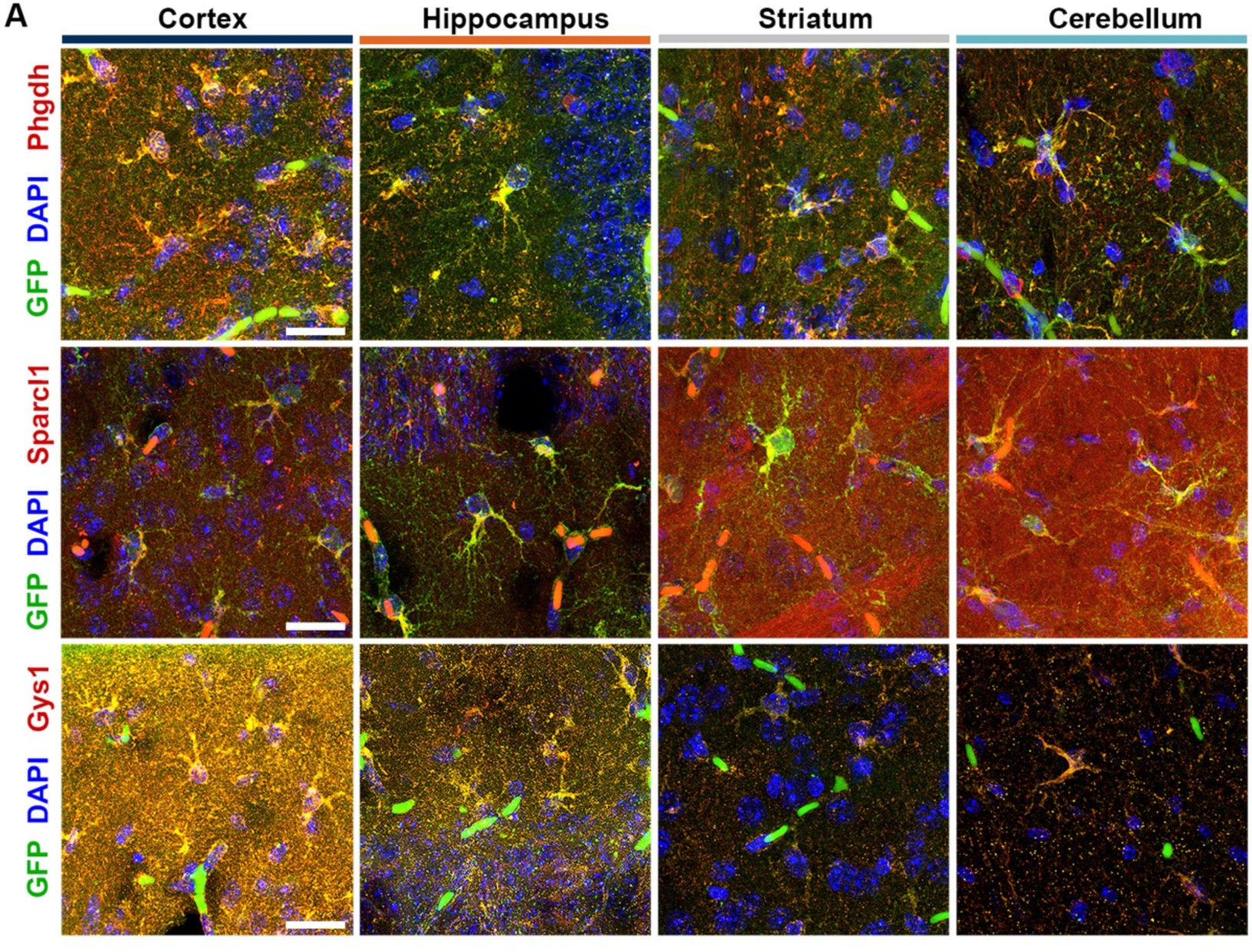
Identification of astrocytic proteins in GFP expressing astrocytes. Representative immunohistochemical stainings of prominent astrocytic proteins. D-3- phosphoglycerate dehydrogenase (*PHGDH*), Sparc-like protein1 (*Sparcl1*) and Glycogen synthase (*Gys-1*) are found in GFP-positive astrocytes of all four brain regions analyzed as expected from identified protein expression pattern by mass spectrometry (Scale=20 µm).

### The cerebellar proteome shows characteristics of Bergmann glia

As proteomic analysis pointed towards stronger differences in the cerebellar dataset, we opted to look deeper into the characteristics of identified proteins in the cerebellum. As two major astrocytic cell types exist in the cerebellum, we performed FUNCAT in cerebellar slices after metabolic labeling to visualize cells that integrated ANL into proteins. Clear TAMRA-positive cells can be observed in the Purkinje cell layer in slices derived from Cre-expressing mice (Cre+) indicating that indeed Bergmann glia, located in the vicinity of Purkinje cell bodies, are the predominantly TAMRA-labeled astrocytes in the cerebellum **(Figure 7A).** In addition, protoplastic astrocytes tagged with TAMRA are also detected within the cerebellar granular layer. This pattern of ANL integration visualized by the TAMRA-tag is consistent with Aldh1l1 expression occurring also in Bergmann glia cells (Doyle et al., 2008).

**Figure 7.**
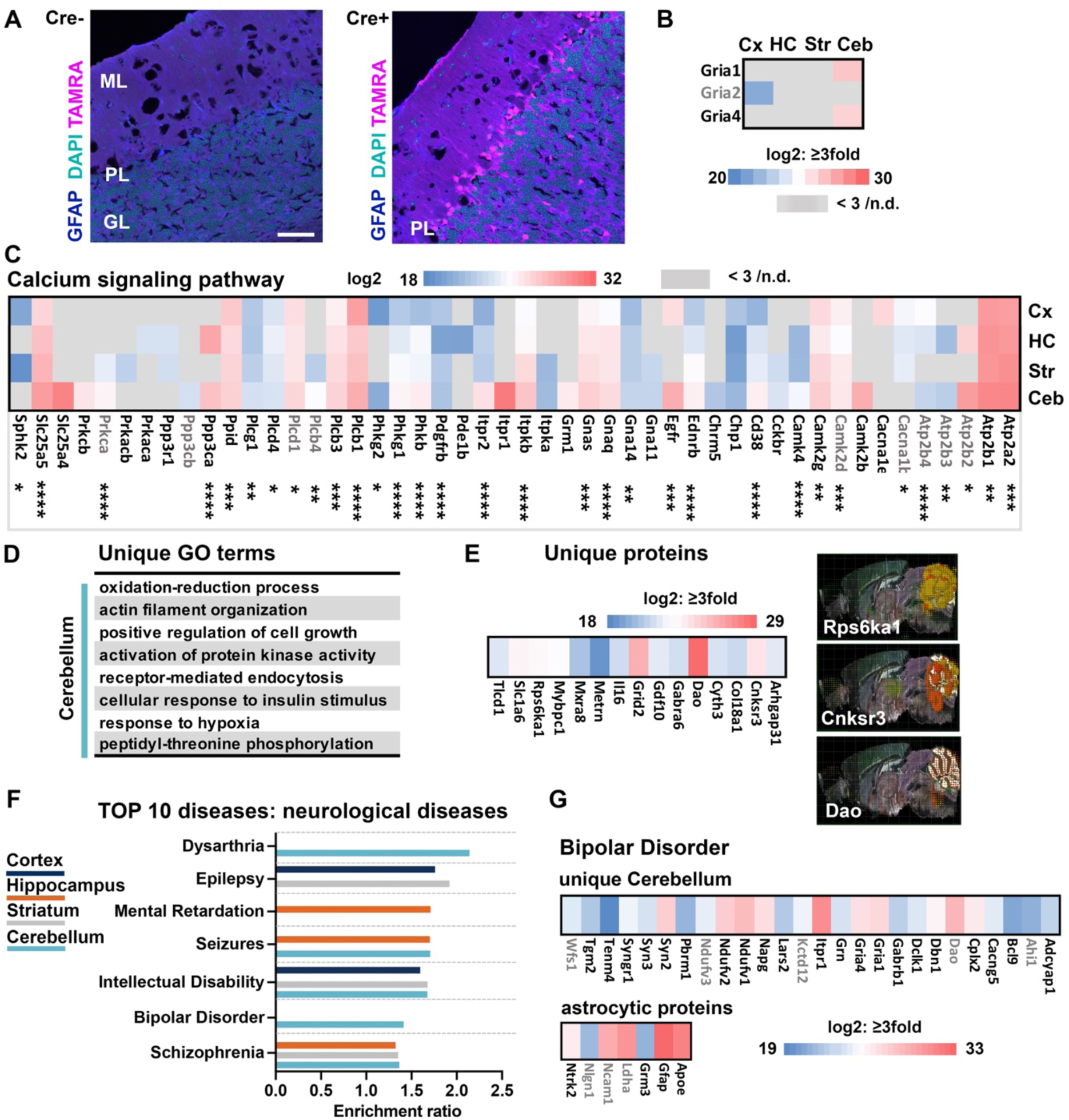
Proteomics of cerebellar astrocytes. **(A)** The TAMRA-tag was visualized after performing FUNCAT in cerebellar slices derived from Cre+MetRS^L274G^ (Cre+) or Cre-MetRS^L274G^ (Cre-) mice upon ANL labeling. TAMRA positive cell somata are found primarily in the Purkinje cell layer (PL) of Cre+ derived tissue. (GL= granular cell layer, ML= molecular layer) Scale=50µm **(B)** Log2 intensities of identified glutamate receptor subunits 1, 2 and 4 (*Gria1,2,4*). *Gria1* and 4 but not *Gria2* are identified in the cerebellum. Gene names in black refer to reviewed protein entries in UniProt. **(C)** All identified and threefold enriched candidates of the KEGG annotation term ‘calcium signaling pathway’ found in the 1D annotation enrichment of annotation terms in Figure 4 are displayed with log2 intensities for each region. Several genes show enhanced log2 intensities in the cerebellum or were only identified in the cerebellum. Gray denotes lack of 3-fold enrichment or detection. One-Way ANOVA was performed to compare abundances (*p<0.05; **: p<0.01; ***p<0.001; ****p<0.0001). Gene names in black refer to reviewed protein entries in UniProt. **(D)** List of GO terms concerning biological processes unique for the cerebellum among the top significant GO biological processes displayed in Figure 4A, B. (see also Supplemental Figure 5A for all GO terms). **(E)** Displayed proteins were exclusively found in the cerebellar dataset, as depicted in the Venn diagram in Figure 2F, and also show highly enriched or unique gene expression pattern in the mouse brain atlas of the Allen Brain Map. Three examples showing in situ hybridization of candidates with high signal in the mouse cerebellum generated by Brain Explorer 2 (“mouse.brain-map.org”). **(F)** Gene enrichment analysis focused on diseases. Over-representation analysis was performed: DisGeNET and WebGestalt (WEB-based GEne SeT AnaLysis Toolkit). The weighted set cover algorithm summarizes each significant term into 10 categories. Only nervous system-related diseases are shown here. **(G)** All 158 gene names annotated to “bipolar disorder” were compared with proteins uniquely identified in the cerebellum or with the list of prominent astrocytic candidate proteins from Figure 5B and plotted with their log2 intensities. (See also Supplemental Figure 7).

Indeed, several characteristic proteins of Bergmann glia are either exclusively found in the cerebellar dataset or highly abundant compared in this brain region. One hallmark of Bergmann glia is the expression of glutamate receptor GluA1 and GluA4 subunits and both subunits are exclusively abundant in the cerebellar dataset **(Figure 7B)**. Looking at the list of key astrocytic proteins **(Figure 5B)**, we find the excitatory amino acid transporter 1 (*Slc1a3*) and the potassium channel Kir4.1 (*Kcnj10*) highly abundant in the cerebellar dataset with both proteins playing relevant roles in neurotransmitter regulation and modulation of synaptic activity by Bergmann glia (De Zeeuw & Hoogland, 2015).

As Bergmann glia are known to respond to neuronal activity with distinctive calcium signaling, we plotted proteins of the calcium signaling pathway based on our 1D annotation enrichment analysis **(see Figure 4D)** with log2 abundances in a heatmap comparing all analyzed brain regions. Indeed, many of the enriched proteins within this pathway show either enhanced abundance or exclusive expression in the cerebellar dataset **(Figure 7C)**. Highly abundant candidates are Inositol 1,4,5- triphosphate receptor type 2 (*Itpr2*), a channel that mediates calcium release from the endoplasmic reticulum or Calcium/calmodulin-dependent protein kinase type II subunit beta (*Camk2b)*.

We further looked into unique processes and unique proteins that might contribute to a specialization of astrocytes within the cerebellum. Biological processes being uniquely enriched in the cerebellum by GO analysis are listed in **Figure 7D**. They include “Receptor-mediated endocytosis” and “Positive regulation of cell growth”, among others. We further screened for proteins that were found uniquely in the cerebellar dataset and also show exclusive or enhanced expression of their mRNA in the cerebellum (“mouse.brain-map.org”; **Figure 7E**). Among them is Meteorin (*Metrn*) that was previously detected in the cerebellum and that plays a role in astrocyte to radial glia differentiation (Nishino et al., 2004). Strong correlation with in situ hybridization data was also found for D-amino-acid oxidase (Dao), an enzyme that regulates D-serine levels and is known to be highly expressed in the Bergmann glia (Horiike et al., 1994), the ribosomal protein S6 kinase α1 (RSK1, Rp6ska1), a S/T protein kinase that acts in the ERK signaling pathway (Romeo et al., 2012) or Cnksr3 that regulates sodium transport and is a negative regulator of ERK signaling (Ziera et al., 2009).

To further elucidate whether astrocytes in different brain regions are associated with nervous system diseases, we performed a gene enrichment analysis focusing on diseases. Indeed, datasets have region-specific disease associations. Here, the hippocampus is specifically associated with “Mental Retardation” **(Figure 7F and Supplemental Figure 7A)**. For the cerebellar dataset we find an exclusive association with “Bipolar disorders” and “Dysarthria”. Dysarthria, a speech sound disorder, and also pathomechanisms of bipolar disorder have been linked to the cerebellum previously (Minichino et al., 2014; Spencer & Slocomb, 2007) but in how far glial cells are involved in disease mechanisms has not been recognized to date.

For “Bipolar Disorder” we found 157 cerebellar proteins annotated to this term. To further identify proteins that specifically contribute to this association, we compared this list with our uniquely identified proteins within the cerebellar dataset. Indeed, several of these candidates are associated with the term “Bipolar Disorder”. Interestingly, a number of these proteins are receptors or channels (*Gabrb1, Gria1, Gria4, Kctd12*) or are mitochondrial proteins (*Ndufv1-3, Lars2*) **(Figure 7G)**. To further understand underlying mechanisms potentially associated with “Bipolar Disorder” we compared the list of disease-associated candidate proteins with our list of key astrocytic proteins with known relevance for neuron-glia interaction **(see Figure 7G, see also Figure 5B).** This indeed identified a couple of proteins, such as Apolipoprotein E (Apoe) or the metabotropic glutamate receptor Grm3 being associated with “Bipolar disorder” **(Figure 7G)**. We further compared 89 genes annotated to the term “Dysarthria” with our cerebellar dataset and identified a number of overlaps **(Supplemental Figure 7B)**.

### Subregional heterogeneity of astrocytic protein translation in the hippocampus

While, probably with the exception of the cerebellum, a comparison between astrocytic regional proteomes revealed rather subtle differences in protein abundance, we sought to get insight into sub-regional diversity of astrocytes. To this end we chose the hippocampal formation with its functionally and structurally well-studied circuitry and anatomy.

Functional diversity as well as differential activation of signaling cascades with consequent cellular adaption processes are reflected by proteome differences and changes in protein expression. To study such differences in astrocytic protein translation in hippocampal subregions, we applied FUNCAT to visualize de novo synthesized proteomes after Cre-dependent activation of MetRS^L274G^ and GFP expression in astrocytes and subsequent ANL integration into newly synthesized proteins. Proteins were visualized *in situ* in hippocampal slices after TAMRA-tagging by ‘click reaction’. Confocal images were analyzed for translational activity utilizing the GFP signal to create a 3D mask for the quantification of the fluorescence-tag within single astrocytes of the hippocampal cornu ammonis (CA)1 and CA3 regions and the dentate gyrus (DG) **(Figure 8A)**. The mean TAMRA intensity, volume and mean GFP intensity of single astrocytes was found to be higher in the CA1 region as compared to CA3 and DG **(Figure 8B-D)**. No significant differences were detectable between DG and CA3 astrocytes. We further analyzed astrocytic translation rates in different layers of the hippocampal CA1, CA3 and DG regions. Here, astrocytes of the CA1 stratum oriens (SO) and to a lesser extend of the CA1 stratum pyramidale (SP) show enhanced mean TAMRA-intensities **(Figure 8E)**. No significant variation of TAMRA intensity was observed in CA3 astrocytes **(Figure 8F)**. In the DG, slightly but significantly enhanced TAMRA intensity was evident in stratum moleculare (SM) astrocytes **(Figure 8G)**. Little variation was seen in GFP reporter expression among groups **(Supplemental Figure 8A-C**). In order to look for potential correlations of differential translational activity with morphological attributes in hippocampal subregions to differentiate possible astrocytic subtypes on a phenotypic basis, we performed Sholl analyses using the 3D mask based on astrocytic GFP expression. Of note, higher morphological complexity of astrocytic process arborization do not correlate with higher translational activity. Astrocytes in CA1 and CA3 display layer-specific differences in their complexity as revealed by Sholl analysis **(Figure 8H, I)**. However, while CA1 astrocytes display similar process lengths **(Supplemental Figure 8D)** astrocytes of the CA3 region reveal layer-dependent differences in overall process length **(Supplemental Figure 8E).** In contrast to CA subregions, astrocytes within DG layers are both homogenous in ramification and total process length **(Figure 8J, Supplemental Figure 8F).**

**Figure 8.**
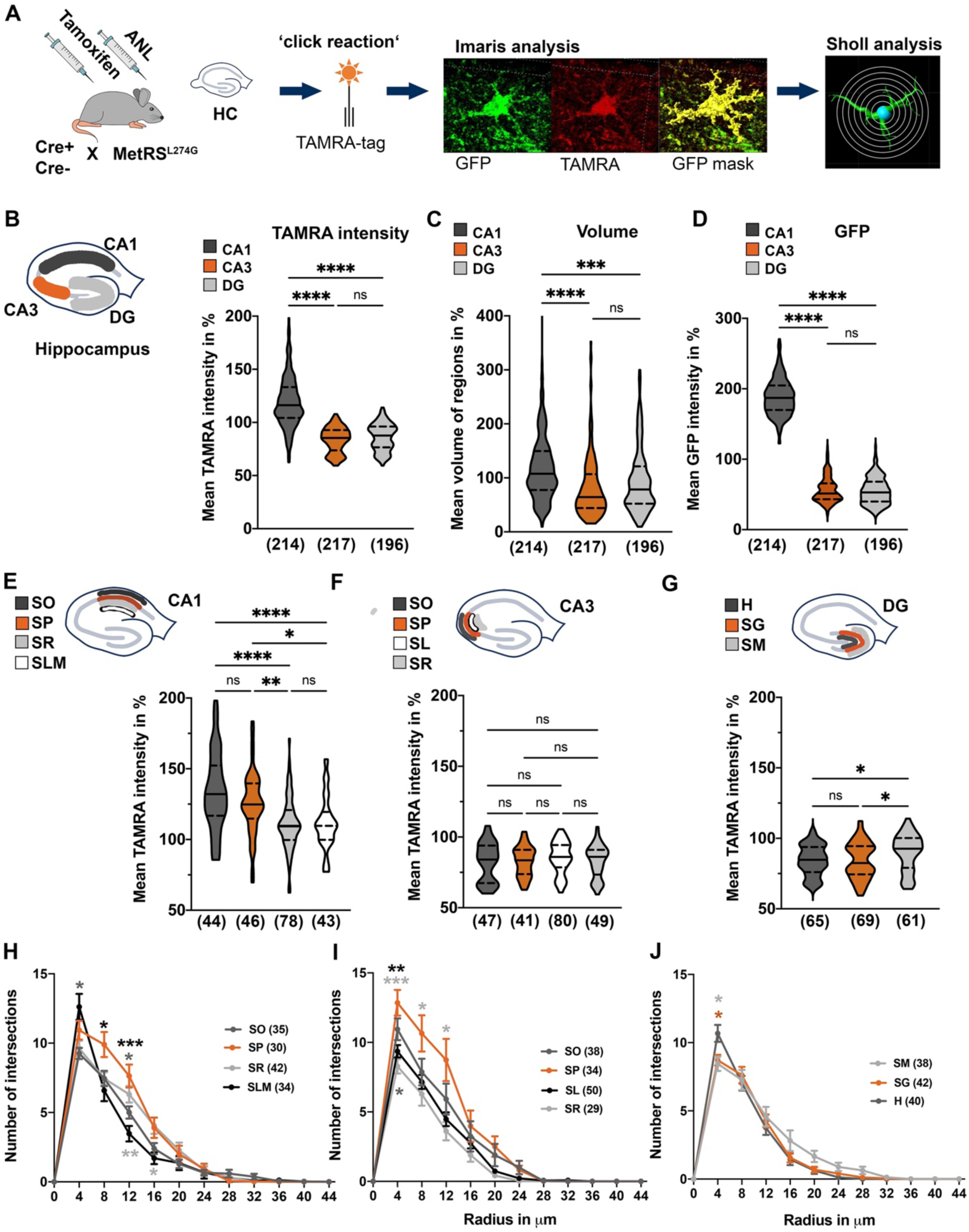
Analysis of astrocytic translation rates in subregions of the hippocampus. **(A)** Working steps for single-cell analysis. Upon Cre recombinase induction and ANL integration, the hippocampus was isolated, sliced and ANL-labeled proteins were tagged with a fluorescence tag (TAMRA-tag) by ‘click reaction’. Imaris (Bitplane Inc.) was used to generate a 3D surface mask of the GFP reporter to determine TAMRA intensity for single astrocytes and to analyze cell morphology or perform Sholl analysis. **(B)** Mean TAMRA intensity of single astrocytes is enhanced in Cornu Ammonis (CA)1 astrocytes whereas CA3 and dentate gyrus (DG) astrocytes show similar levels. (lines indicate median and upper/ lower quartiles; One-Way ANOVA; F(2)=288.2, p<0.0001; multiple comparisons: ****p<0.0001;). **(C-D)** CA1 astrocytes show increased mean volumes **(C)** and GFP expression **(D)** (One-Way ANOVA; C: F(2)=2745; p<0.0001, D: F(2, 623)=21.09, p<0.0001), multiple comparisons; ****p<0.0001; ***p<0.001;. **(E-G)** Mean TAMRA intensity of astrocytes from sublayers of CA1 **(E)**, CA3 **(F)**, DG **(G)** were determined. **(E)** In CA1 astrocytes of in the stratum oriens (SO) or in the stratum pyramidale (SP) show strongest TAMRA intensities. (SR: stratum radiatum; SLM: stratum lacumosum moleculare) **(F)** No major differences were observed in CA3 (SL: stratum lacumosum) **(G)** Increased TAMRA intensity is observed in stratum moleculare (SM) of DG (SG: stratum granulosum; H: hilus; One-Way ANOVA; E: F(3)=13.51, p<0.0001; G: F(2)= 4.591, p=0.0113; multiple comparison: *p<0.05; **p<0.01; ****p<0.0001;) **(H-J)** Sholl analysis of individual astrocytes of sublayers of hippocampal CA1**(H)**, CA3 **(I)** or DG areas **(J; T**wo-way ANOVA, region factor: H: F(3)=2.558, p=0.0577; I: F(3)=5.307, p=0.0017; J: F(2)=0.6209, p=0.5392; multiple comparison: *p<0.05; **: p<0.01; ***p<0.001). Astrocytes of CA1 and CA3 SP show elaborated process ramification. (Numbers in brackets indicate number of cells used for analysis. For further analyses see Supplemental Figure 8).

This first peek into subregional astrocyte diversity at the level of translational activity and general morphological features further promotes the idea of subregional and circuit specificity of astrocytes in the hippocampal formation.

## Discussion

There is mounting evidence that astrocytes are a diverse group of cells that show inter- and intraregional specializations (Khakh & Deneen, 2019). Here, we established a mouse line that enables astrocyte-specific *in vivo* labeling of proteins utilizing the induced expression of the modified methionyl-tRNA synthetase MetRS^L274G^ and the methionine surrogate ANL, which allows subsequent isolation and identification of tagged proteins to study regional proteome differences in astrocytes or visualization of general translation activity in single astrocytes (Alvarez-Castelao et al., 2017; Erdmann et al., 2015; Müller et al., 2015). Evaluating the specificity of MetRS^L274G^ and GFP expression upon Cre recombinase induction by tamoxifen revealed a high cell-type specificity: the GFP reporter and tagged proteins were only visible after induction by tamoxifen, they were restricted to astrocytes and were comparable among brain areas. LC-MS/MS data obtained from four different brain areas further support astrocyte specificity. The data overlap with previously identified astrocytic proteins, either with published proteomic data from Sharma et al. (Sharma et al., 2015) or with AstroProt, a database we created as part of this study.

We performed GO and annotation enrichment analyses which both disclose a prevalence for pathways and functions being involved in metabolic processes. This is well in line with recent finding that astrocytes play a prominent role in regulating molecular and metabolic homeostasis of neurons (Verkhratsky & Nedergaard, 2018). A similar prevalence of metabolic processes reflected by the astrocytic transcriptome/proteome has also been shown by other studies (Chai et al., 2017; Ohlig et al., 2021) further supporting the specificity and reliability of the present approach as a versatile tool to analyze astrocyte-specific proteomes in diverse brain regions with high cell-type specificity.

A comparison of datasets obtained from all four brain regions reveals many commonalities between astrocytes of different brain regions. Both gene ontology (GO) and cluster analysis indicate that several biological processes are common or shared by at least two regions. Nevertheless, significantly enriched GO terms being exclusive for single brain regions are among identified processes and might reveal regional astrocytic specialization.

In line with process analysis, we see a clear overlap of proteins shared by all datasets supporting findings from Chai *et al*. who found a strong overlap of hippocampal and striatal astrocytic proteomes (Chai et al., 2017). We observe differences in protein abundances but only a minor fraction of proteins has regionally restricted expression pattern. In how far region-specific abundances of certain pathways or proteins impact on regional differences in astrocytic function and neuronal support is one of the major questions and is yet to be determined in detail.

In agreement with previous transcriptomic studies that were able to identify changes in gene expression pattern but also observed major differences of astrocytic morphology, physiology and molecular function with subregional resolution (Bayraktar et al., 2020; Farhy-Tselnicker et al., 2021; Lanjakornsiripan et al., 2018; Lin et al., 2017; Morel et al., 2017), we find differences in general translation rates of astrocytes in subregions of the hippocampus among CA1, CA3 and DG astrocytes as well as within CA layers, supporting the idea that indeed astrocytes have differential subregional and layer-specific dynamics of protein synthesis-dependent cellular processes.

By bioinformatic analyses, we found a clear distinction of the astrocyte-specific cerebellar dataset compared to other datasets analyzed. Cerebellar processes are highly prominent in the KEGG-based annotation enrichment analysis and also cerebellar proteins show a clear difference in clustering and protein abundance. By using labeling with a TAMRA tag, we find many astrocytes labeled in the Purkinje layer of the cerebellum likely representing Bergmann glia, proving that these cells strongly contribute to the isolated cerebellar proteome. These Bergmann glia present a subgroup of astrocytes with distinct morphological and physiological features (Buffo & Rossi, 2013) but little is known about their proteomic composition and a general characterization, to date, is restricted to transcriptomic data (Koirala & Corfas, 2010).

Also, proteomic data point towards Bergmann glia as a source for tagged proteins. We note a predominant abundance of the glutamate transporter GLAST-1 (*Slc1a3*), which is, given its strong expression, often used as marker for Bergmann glia cells in the cerebellum. We further identify abundant candidates relevant for the support of synaptic signaling such as the potassium transporter Kir4.1 (*Kcnj10*) and both glutamate receptor subunits 1 and 4 (*Gria1, 4*), previously described to mediate calcium entry into Bergmann glia (Saab et al., 2012).

Given its prominent role in Bergmann glia, we sought to further identify proteins relevant in the “Calcium signaling pathway”, one of the exclusively enriched processes in the cerebellum applying the KEGG-based analysis. This revealed several candidates that are highly enriched in the cerebellar dataset, including InsP3R2 (*Itpr2*) with its well know contribution to astrocytic calcium signaling (Holtzclaw et al., 2002; Srinivasan et al., 2015). Other abundant candidates such as the ADP/ATP translocase 1 and 2 (*Slc25a4, 5*) were not linked to astrocytic calcium signaling in the nervous system, so far.

A look at disease associations by GO analysis, revealed regional differences, with a strong association of hippocampal astrocytic proteins to “Mental retardation” and cerebellar proteins to “Dysarthria” and “Bipolar disorder”. Annotated genes for “Bipolar disorder” reveal an association with synapse regulation and function as well as mitochondrial contribution. In fact, a role of the cerebellum in mood disorders, including the bipolar disorder, was hypothesized previously, largely based on neuroimaging studies (Minichino et al., 2014). Initial insights into the molecular basis of this disease involves also mitochondrial complex I subunits, that were also identified in our dataset. Changes in expression of complex I proteins was associated with mood disorders and a reduction of expression in the cerebellum was identified in bipolar disorder patients (Ben-Shachar & Karry, 2008). In how far astrocytes contribute to onset and/or disease progression of mood disorders in the cerebellum remains elusive. However, astrocytes were previously associated with bipolar disorder and several molecular targets affecting also astrocytic function are in the focus as treatment targets for mood disorders (Di Benedetto & Rupprecht, 2013; Keshavarz, 2017).

With this study we proof the applicability of this method to investigate astrocytic proteomes in different areas of the mouse brain. Whereas highly sensitive transcriptomic studies can provide detailed information about regional and cellular differences in gene expression patterns of astrocytes (Batiuk et al., 2020; Bayraktar et al., 2020; Zeisel et al., 2015), it is much more difficult to get information about the actual proteome in these cells with similar resolution and sensitivity. Yet, this information is important as studies comparing transcriptomic and proteomic data show that mRNA and protein abundance do not necessarily correlate completely (Chai et al., 2017; D. Wang et al., 2019; Zhang et al., 2014). Hence, proteomic studies in combination with other omics techniques are needed to fully characterize the molecular composition and regional specialization of astrocytes. The strategy established here can contribute to such a multi-omics approach.

Although with our labeling technology a bias towards overrepresenting proteins with a high turnover cannot be excluded, nonetheless, as most proteins have a half-life time between 3-7 days (Fornasiero et al., 2018), we likely are able to identify a broad range of proteins with the applied labeling time frame. Indeed, comparing our cortex dataset with a dataset obtained after astrocyte isolation (Sharma et al., 2015), we cover a similar spectrum of proteins, implying that our datasets reflect an unbiased portion of the astrocytic proteome.

One major advantage of this technique is the *in vivo* nature of labeling that ultimately allows analysis of proteome changes in diverse steady state conditions or disease conditions. Due to the *in vivo* nature, a combination of this tool with behavioral paradigms using variable labeling duration and application modes of ANL, will be of particular interest for future questions regarding the role of astrocytic physiological adaptations and their contribution to behavior as has been shown for the neuronal proteome (Alvarez-Castelao et al., 2017). With this technique and it’s *in vivo* application in astrocytes, we aim to contribute to a further characterization of astrocyte diversity and function in health and disease.

## Methods

### Animals

The present study was carried out in accordance with the recommendations of the National Committee for the Protection of Animals Used for Scientific Purposes of the Federal Republic of Germany and European regulations for ethical care and use of laboratory animals (2010/63/EU). The experiments were approved by the local Ethics Commission of the Federal State of Saxony-Anhalt (42502-2-1329 Uni MD). The animals were housed under controlled laboratory conditions at constant temperature (20±2°C) and air humidity (55-60 %). 12h light-dark cycles were used (lights on at 6:00 a.m.). Animals had access to commercial food pellets and tap water *ad libitum*. The health status of animals was monitored regularly.

Homozygous (fl/fl) or heterozygous (fl/+) mice (*Mus musculus*) of the line MetRS^L274G^ (also MetRS*; C57BL/6-Gt(ROSA)26Sor^tm1(CAG-GFP,-Mars*L274G)Esm^/J, Jax.org stock nr: 028071) (Alvarez-Castelao et al., 2017) were used and crossed with a Cre-recombinase mouse driver line Aldh1l1-CreERT2 (Winchenbach et al., 2016). Adult animals, positive (Cre+) or negative (Cre-) for Cre-recombinase coding sequence and for MetRS^L274G; fl/fl^ or MetRS^L274G; fl/+^ were used for initial testing of labeling efficiency (see Figure 1). Further experiments were performed with 8-12-week-old mice MetRS^L274G; fl/fl^ of both sexes.

### *In vivo* metabolic labeling

Tamoxifen (Merck KGaA, Darmstadt, Deutschland, T5648) was applied as described (Winchenbach et al., 2016). In brief, tamoxifen was dissolved in corn oil (Merck, C8267) at 7,5 mg/ml and administered intraperitoneally (i.p.; 75 μg/g body weight) once a day for 5 consecutive days. After a break of 9 days, Azidonorleucine (ANL; synthesized according to Link et al., 2004) or methionine (Met; Merck, M5308), dissolved in 0,9% NaCl solution (NSS Serumwerk Bernburg, Germany), were administered i.p. (2 μmol/g body weight) once a day for 4 consecutive days. Cycles of 24 hours were respected between subsequent i.p. injections.

### Tissue preparation

The animals were sacrificed by neck dislocation and hippocampus, striatum, cortex and cerebellum were extracted and fresh frozen for further mass spectrometry or biochemical analysis. For FUNCAT and morphometric analysis, 450 µm thick hippocampal slices were trimmed with a tissue chopper (The Mickle Laboratory Engineering Co. LTD, United Kingdom; MTC/2E McIlwain Tissue Chopper) or whole mouse brains were sliced coronally for the regional analysis using Mouse brain blocker (PA002, David Kopf Instruments, Tujunga, CA, USA). Slices were quickly fixed overnight (ON) in 1x PBS pH7.4, 4% sucrose (Serva, Heidelberg, Germany) and 4% PFA (Paraformaldehyde; Thermo Fisher Scientific, Waltham, USA). Fixed brain slices were washed once with 1x PBS pH 7.4 and subsequently submerged in 30% (w/v) sucrose for two days and then embedded in tissue freezing medium (Leica Microsystems, Milton Keynes, United Kingdom; 14020108926).

### Fluorescent tagging (FUNCAT)

The FUNCAT protocol was done according to Dieterich *et al*. Dieterich et al., 2010 with some modifications. In brief, hippocampal slices were resliced (20 μm) with a cryostat (Leica Microsystems, Milton Keynes, United Kingdom, CM1950) and collected in 1x PBS pH 7.4. The slices were treated with 0.5% Triton X-100 in blocking solution (10% (v/v) horse serum (Thermo Fisher Scientific), 5% (w/v) sucrose, 2% bovine serum albumin (BSA; v/v) in 1x PBS pH 7.4) and incubated ON at 4°C (from now on, all steps were done under gentle agitation, 30±5 rpm). The slices were washed thrice for 10 min each at room temperature (RT) with 1x PBS pH 7.8 with 0.1% Triton X-100, followed by three washing steps for 5 min, RT each with 1x PBS pH 7.8. The FUNCAT reaction consists of 40 μM Triazole ligand (Tris(1-benzyl-1H-1,2,3-triazol-4-yl)-methylamine; TBTA; Merck; 678937), 0.1 mM TCEP (tris-(2-carboxyethyl)phosphine; Merck, C4706), 0.4 μM 5/6- TAMRA-PEG 4 -Alkyne (Jena Bioscience; Jena, Germany; CLK-TA108) and 0.2 mM freshly prepared CuSO_4_ in 1x PBS pH7.8, as described in detail previously (Dieck et al., 2012). The slices were incubated ON at RT in the FUNCAT reaction mixture in the dark. Following day, slices were washed thrice for 20 min at RT each with wash buffer (0.5 mM EDTA, and 1% Tween-20 (v/v) in 1x PBS pH 7.4) and twice for 10 min at RT each with PBS pH 7.4 containing 0.1% Tween-20 (v/v). Further immunohistochemistry staining was performed with treated slices.

### Immunohistochemistry

For immunohistochemistry, all steps were performed under gentle agitation (30 ± 5 rpm) and protected from light. The slices were washed thrice 5 min at RT with TPBS (0.12% Tris, 0.9% NaCl, 0.025% NaH_2_PO_4_) and incubated with primary antibodies dissolved in TPBS-TS-1% (TPBS added with 1% Horse serum and 0.25% Triton X-100). Slices were stained with anti-GFAP (Abcam, Cambridge, United Kingdoms; ab4674) or anti-MAP2 (Synaptic Systems, Goettingen, Germany; 188004) and anti-GFP (Sicgen Antibodies, Cantanhede, Portugal; AB0020) and TAMRA fluorescence tag was intensified with an Anti-TAMRA antibody (Thermo Fisher Scientific; MA1-041) and incubated ON at 4°C. Slices were washed four times with TPBS-T (TPBS with 0.3% Triton X-100) for 5 min each at RT and incubated with secondary antibodies in TPBS-TS-1% for 4h at 4°C (Alexa Fluor 546 anti-Mouse (Merck, A-11018), Alexa Fluor 488 anti-goat (Abcam; ab150129) and Alexa Fluor 647 anti-chicken (Jackson Immunoresearch Europe Ltd.Cambridgeshire, United Kingdom; 703-606-155). Slices were washed thrice with TPBS at RT for 5 min each and incubated with DAPI at RT for 15 min, washed thrice with TPB for 5 min at RT and mounted in Mowiol (Carl Roth, Karlsruhe, Germany). The same protocol was used for staining of candidate proteins omitting steps for FUNCAT reaction. Coronal whole brain slices (20µm) were incubated with primary antibody (Anti-PHGDH (3-Phosphoglycerate dehydrogenase; Atlas antibodies, Bromma, Sweden; HPA021241), Anti-SparcL1(Sparc-Like1; R&D systems, Minneapolis, USA; AF2836), Anti-Gys1 (Glycogen Synthase1; Proteintech Group, Rosemont, USA; 10566-1-AP) directly after blocking.

### Microscopy and image processing

Immunohistochemistry staining in combination with FUNCAT were analyzed via confocal laser scanning microscopy (CLSM; Axio observer.Z1 microscope provided with a LSM 710 confocal system; Carl Zeiss group) was used together with its user interface ZEN 2012 SP5 (Carl Zeiss, Oberkochen, Germany). Cells for all analyses were obtained from six mice with uniform distribution. All images were taken in Z-stacks with an optical step of 0.45 µm and in 16-bit 4x4 tiles0.08 µm pixel size under Plan-Apochromat 63x/ numerical aperture 1.4 oil DIC M27 immersion objective. Images were stitched using image-processing built-in software. Identical exposure times for all hippocampal slices and regions analyzed were used. Microscopy settings remained the same in all images scanned for GFP and TAMRA channels.

The z-stacked images were imported into IMARIS software of Version 9.5/9.6 (Bitplane AG, Zürich, Switzerland) to quantify the intensities in 3D astrocyte structures. The 3D mask of astrocytes was created with the GFP signal using the IMARIS surface module and the continuous astrocyte surface that show a distinct soma in the center are used as 3D masks for quantifying the TAMRA, GFP intensity and volume for each cell. Mean TAMRA, GFP intensity or volume values in subregions of the hippocampus or different hippocampal layers were related to the respective mean intensity or volume of total number of analyzed hippocampal astrocytes. The morphology of astrocytes was quantified using ‘Filament tracer’ of IMARIS with Gaussian filter set to 0.25 µm. This module creates an autopath with starting diameter point of 4 µm and seed point of 0.176 µm to reach until the edges of all branches of astrocytes. Missing branches were set manually or automatically set seed points within the cell soma were manually removed. As a result, the data for the number of 3D Sholl intersections and sum of all branch length were procured from this analysis.

### Bioorthogonal non-canonical amino acid tagging (BONCAT)

Brain tissues were homogenized by Potter S (Sartorius, Goettingen, Germany) in 1x PBS (10ml/1g), pH 7.8, containing EDTA-free cOmplete™protease inhibitor cocktail (PBS-PI; Roche, Basel, Schweiz) and 250 U/ml Benzonase®nuclease (Merck).

The BONCAT procedure was performed as essentially described with minor modifications (Dieterich et al., 2007; Landgraf et al., 2015; Müller et al., 2015). In brief: Homogenates were solubilized with SDS to a final concentration of 0.1% and boiled at 95°C for 5 min in 25% of final volume. To cooled samples, 0.2% TritonX100 and 1x PBS-PI was added to reach final volume. The samples were centrifuged for 5 min at 3000 g, 4°C. Supernatants underwent click-chemistry. For that, protein extracts were adjusted to 0,5 µg/ml concentration and sequentially supplemented with 0.2 mM triazole ligand (Tris[(1-benzyl-1H-1,2,3-triazol-4-yl)methyl]amine; TBTA, Merck; 678937), 25 µM biotin-alkyne tag and 0.2 mg/ml copper(I)bromide-suspension with 10 sec vortexing steps in between steps. Samples were incubated 90 min at RT using an end-over-end mixer. Samples were centrifuged at 3000 g, 4°C for 5 min and the supernatant further processed for immunoblotting.

### Immunoblotting

The supernatant collected after BONCAT (1.1.8) was either boiled with 4x SDS sample buffer (4% SDS, 40% glycerol, 20% ß-Mercaptoethanol in 250 mM Tris-HCl, Merck) at 95°C, 5 min for biotin detection or 70°C, 15 min for GFP quantification. The samples were run on 5-20% SDS-polyacrylamide gels and transferred or spotted onto nitrocellulose membrane with 2µg/µl concentration using Bio-dot® Microfiltration Apparatus (Bio-Rad, Feldkirchen, Germany). Total protein amounts per lane were stained with No-stain™ Protein Labelling Reagent (Thermo Fisher Scientific; A44449) and used for lane or dot normalization. At least two technical replicates were done for all quantifications. Primary antibodies were used: 1:10000 rabbit anti-biotin (Fortis Life Sciences, Waltham, USA, A150-109A), and 1:1000 rabbit anti-GFP (Thermo Fisher Scientific, A-11122). Secondary antibodies were peroxidase-conjugated donkey anti-rabbit IgG (Dianova, 711-035-152) for biotin detection or anti-rabbit IR Dye 800nm (Li-COR, Lincoln USA; 827-08365) for GFP detection and for dot blot analysis. All Imaging and quantifications were done using Image Studio™ software and LICOR Odyssey^FC^ (LI-COR).

### Affinity purification of ANL-labeled proteins for MS analysis

ANL-labelled proteins were affinity purified using the Click-IT® Protein Enrichment Kit (Thermo Fisher Scientific) according to the manufacturers protocol. The protein concentration of each lysate was adjusted to 8mg/ml using the provided lysis buffer and the azide-bearing proteins were coupled via click reaction to an alkyne agarose resin. Afterwards, the samples were reduced using 10 µM DTT, followed by alkylation with 40 mM iodoacetamide solution. Subsequently, resins were extensively washed consistently according to the kit protocol and the bound proteins finally digested using 0.1 µg/µl Trypsin. The resulting digests were desalted using C-18 cartridges and the eluates finally dried in a vacuum concentrator.

### LC-MS/MS Analysis

LC-MS/MS analyses of purified and desalted peptides were performed on a Dionex UltiMate 3000 n-RSLC system (Thermo Fisher Scientific), connected to an Orbitrap Fusion^TM^ Tribrid^TM^ mass spectrometer (Thermo Fisher Scientific). Peptides of each sample were loaded onto a C18 precolumn (3 μm RP18 beads, Acclaim, 0.075 mm × 20 mm), washed for 3 min at a flow rate of 6 µl/min, and separated on a 2m Pharmafluidics C18 analytical column at a flow rate of 300 nl/min via a linear 60 min gradient from 97% MS buffer A (0.1% FA) to 32% MS buffer B (0.1% FA, 80% ACN), followed by a 30 min gradient from 32% MS buffer B to 62% MS buffer B. The LC system was operated with the Thermo Scientific SII software embedded in the Xcalibur software suite (version 4.3.73.11, Thermo Fisher Scientific). The effluent was electro-sprayed by a stainless-steel emitter Thermo Fisher Scientific). Using the Xcalibur software, the mass spectrometer was controlled and operated in the “top speed” mode, allowing the automatic selection of as many doubly and triply charged peptides in a 3-s time window as possible, and the subsequent fragmentation of these peptides. Peptide fragmentation was carried out using the higher energy collisional dissociation mode and peptides were measured in the ion trap (HCD/IT).

MS/MS raw data files were processed via Proteome Discoverer 2.4 mediated searches against the murine UniProtKB/SwissProt protein database (release 2020_01) using Sequest HT as search machine. The following search parameters were used: enzyme, trypsin; maximum missed cleavages, 1; fixed modification, carbamidomethylation (C); variable modification, oxidation (M); peptide tolerance, 7 ppm; MS/MS tolerance, 0.4 Da. The FDR was set to 1%.

### Bioinformatics

#### General

Further processing and analysis of data obtained from Sequest HT as search machine was done using Excel, PANTHER (http://www.pantherdb.org/), Cytoscape/ClueGO (Bindea et al., 2009), Perseus (Tyanova et al., 2016) and inhouse-scripts (written in R and Perl). To obtain a list of specifically enriched proteins, the calculation of the fold change was performed by dividing the average abundance of each sample (i.e average of biological and technical replicates) by the average abundance of the region-specific control obtained from Cre- mice. We defined a protein with a fold change ≥ 3.0 as specifically enriched. All proteins absent from Cre- control but at least present in one sample, were added to the list of enriched proteins.

Cluster analysis was performed in Perseus with log2 transformed abundances and the average log2 intensities of 3-fold enriched proteins were displayed. The top 47 proteins were extracted from the top clusters after the complete cluster analysis **(Supplemental Figure 6A)**.

Single protein candidate comparison for either identified genes of the calcium signaling pathway obtained by 1D annotation enrichment or selected astrocytic candidates is based on average log2 intensities of 3-fold enriched proteins.

#### Validation of data

The cell-specific marker proteins were obtained by determination of 3-fold cell-specific abundance of a previous study determining a cell-type resolved mouse proteome(Sharma et al., 2015). Calculation of the 3-fold abundance results in 713 astroglial and 677 neuronal marker. These were compared with our region-specific datasets.

#### Outlier detection

To detect outliers, we plotted the region-specific log2-transformed abundances of each sample to each other sample or control **(Supplemental Figure 3)**.

#### Volcano Plots

For pairwise comparison of brain regions, -log10 transformed p-values were calculated in Perseus and plotted against log2 intensities. Single candidates with high significance and high difference to the compared dataset were displayed with their respective gene names **(Supplemental Figure 6B)**.

#### Protein classes

For a functional analysis of protein classes we used PANTHER (http://www.pantherdb.org/) (Thomas et al., 2003). We compared glial marker proteins from Sharma et al. 2015 with specifically enriched proteins from the cortex. Only the top 10 protein classes (according to their number of proteins) were visualized.

#### Cytoscape/ClueGO

For the gene enrichment analysis in Fig. 3C Cytoscape 3.8.2 with the Plugin ClueGO 2.5.7 (incl. Cluepedia) was used. Only annotations from gene ontology Biological Process (BiologicalProcess-EBI-Uniprot-GOA-ACAP-ARAP (13.05.2021)) with experimental evidence were applied. Following parameters determined calculation of the significance and visualization: no GO-Term fusion; p<=0.001; 50 Min # genes; only enriched annotation terms; ClueGO-Layout: Repulsion factor: medium; Edge length factor: maximal. Small labels were hidden, and clusters were rearranged to improve clarity.

#### 1D annotation enrichment

The algorithm of the one-dimensional annotation enrichment analysis uses the absolute intensities of each enriched protein as input (≥ 3fold change) and the annotation database KEGG (Kyoto Encyclopedia of Genes and Genomes, https://www.genome.jp/kegg/) and is a part of the Perseus software(Cox & Mann, 2012; Tyanova et al., 2016). For each annotation term a position score is calculated by comparing the corresponding logarithmized protein intensities with the global distribution of all values of all proteins. The significance of the log protein intensities of an annotation term being systematically larger than the global distribution, a two-sided Wilcoxon-Mann-Whitney test was performed. After an FDR correction (Benjamini & Hochberg, 1995) (threshold 0.003), a position score between -1 and 1 was calculated for each significantly enriched annotation term. We generated an annotation matrix of all four regions and clustered all position scores with Perseus.

#### Diseases

For the analysis of significantly enriched pathways involved in diseases we used WebGestalt (WEB-based GEne SeT AnaLysis Toolkit, http://www.webgestalt.org/) (J. Wang et al., 2013). The over-representation analysis was performed with the disease database DisGeNET (https://www.disgenet.org/), minimum number of genes were 30, a FDR correction was done with a threshold of 0.05. The weighted set cover algorithm summarizes each significant term into 10 categories **(Figure 7F, Supplemental Figure 7A)**.

#### Database comparison

The comparison of area-specific proteomes with astroglial- and synapse-specific databases is based on a simple overlap ratio which is calculated by the number of common proteins divided by unique proteins. In this study, the four area-specific protein lists were compared to a) 1,018 proteins from SynGO (https://syngoportal.org/; (Koopmans et al., 2019), b) 2,316 proteins from SynProt (https://www.synprot.de/) (Pielot et al., 2012), c) 9,705 proteins from AstroProt (https://www.synprot.de/) and, to exclude statistical effects due to different sample sizes, to two random protein lists: d) 1,044 proteins (Random 1) and 9,659 proteins (Random 2). All proteins in all comparisons were from the mouse proteome.

#### Gene enrichment analysis (inhouse scripts)

The Inhouse scripts for the singular enrichment analysis (**Figure 4 A, B**) were realized in Perl and R as decribed previously (Lang et al., 2018). The algorithm compares the annotations using the gene ontology database (only Biological Process, 2020/26/09) for each region with 10 different random protein lists of approximately the same size. The random lists were generated by a simple script using the mouse proteome from UniProt (2020/26/09). Only if a certain annotation was found significantly enriched (Fisher’s exact test) after FDR correction (Benjamini & Hochberg, 1995) in all 10 comparisons, this annotation was judged as robustly enriched in the region.

### AstroProt

For a comprehensive analysis we collected data from proteomic and selected transcriptomic studies dedicated to astrocytes (publication dated ≤2018). For this, publications were selected to extract lists of proteins or genes and, in addition, further information about methods, fractions and ages of cells, if available. A database was designed to incorporate all data and made publicly available for academic purposes within the framework of the SynProt-portal (www.synprot.de).

### Statistical analysis

Statistical analysis for quantification of immunoblotting, single cell intensities and morphological data were run in GraphPad Prism 8 and data were presented as bar diagrams with mean±SEM (standard error mean). Two-tailed t-test or One-way ANOVA followed by Turkey test was performed for blot quantification, intensities, volume and branch parameters of single astrocytes. Repeated measurement Two-way ANOVA followed by Holm-Šídák test was conducted for Sholl analysis of astrocytes. Results of two-way ANOVA were displayed as line plot with mean ± SEM for every radius. The number of cells analyzed for statistics is presented under each graph and the number of animals is stated under each figure’s caption. Statistical significance was declared as P-value ≤0.05 for all the datasets and every * represents the times of p-value is less than 0.05.

Statistical comparison of protein abundances were done by t-tests or One-way ANOVA in Perseus. Quantification for p-values ≤0.05 were considered as significant.

## Acknowledgements

This work was supported by DFG SPP1757 and SPP1172. P.P. is associated with RTG 2413 Synage. A.M. and R.P. are supported by CBBS (A.M, RP: EFRE:ZS/2016/04/78113).

We thank Erin Schuman for providing the transgenic mouse line R26-GFP-2A-MetRS^L274G^. Thanks to the Leibniz institute for Neurobiology and to Dr. O. Kobler for providing the Imaris platform and the support in imaging and software usage. We thank José Vázquez López for first technical establishments and we thank Michaela Böx and Grit Borkhardt for the excellent technical assistance and Kathrin Freke for animal care taking.

## Author Contributions

P.P. established labeling methods and performed mouse injections, tissue preparation, FUNCAT, immunohistochemistry, confocal microscopy, immunoblotting, image analysis, data analysis, statistics. R.P. performed bioinformatic analysis, established AstroProt. P.L. established affinity purification, sample preparation, supported bioinformatics, synthesized ANL and alkyne-biotin-tag. A.B. Performed immunoblotting and generated transgenic mouse lines. J.W. und M.v.H. established protein detection and performed LC-MS/MS. E.G. supported AstroProt generation. L.J. supervised sample detection by LC-MS/MS. D.C.D. supervised the project and acquired funding. A.M. supervised the project, performed data analysis, prepared figures, wrote the manuscript with input from P.P., R.P. and P.L. with further contribution from all authors.

## Competing Interests

The authors declare no competing interests.

## Supplemental Figure Legends

**Supplemental Figure 1.**
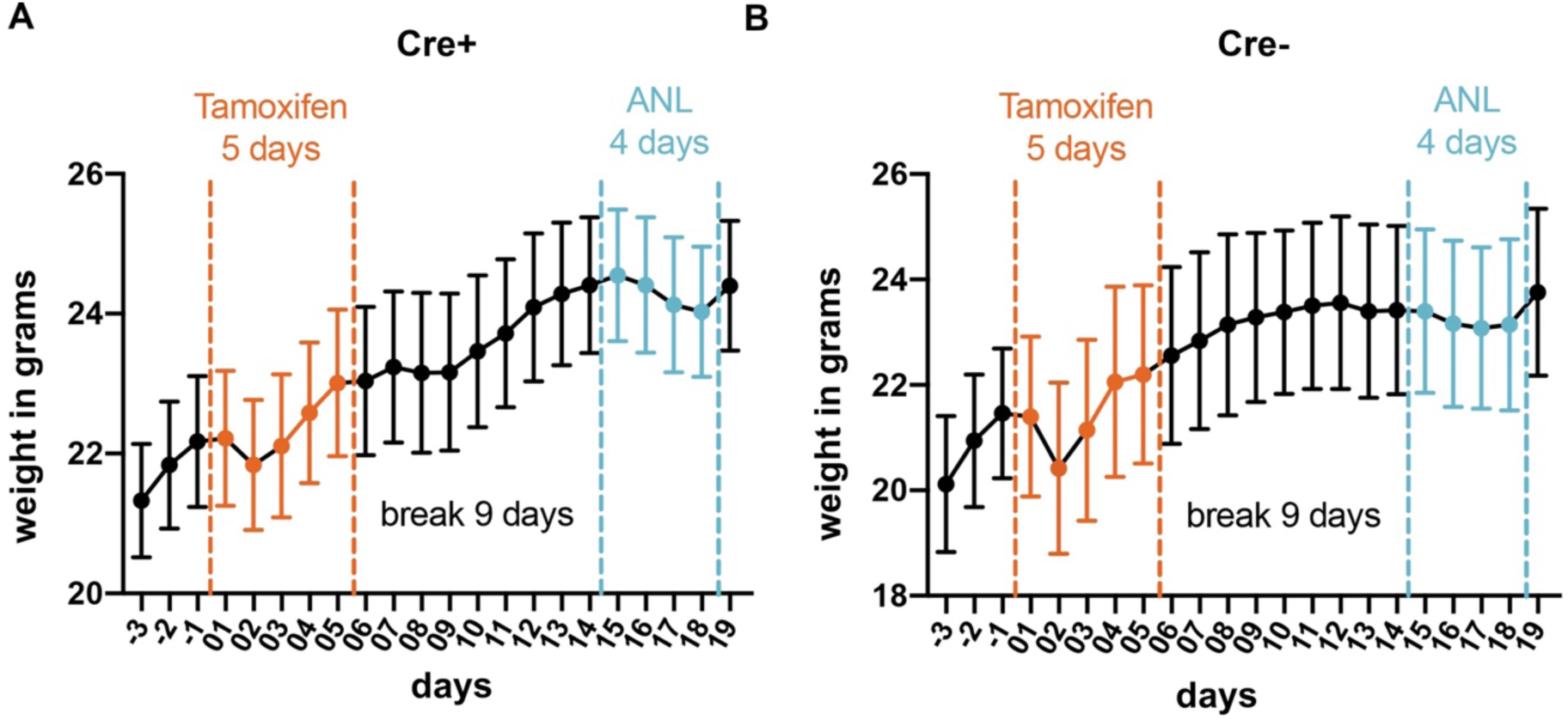
Weight control. (A) Weight control for Cre recombinase expressing MetRS^L274G^ (n=11) mice or Cre- control mice (n=5; mean ± SEM). Slight weight reductions occur with start of i.p. injections with tamoxifen or ANL for Cre+ as well as for Cre- mice. (Related to Figure 1).

**Supplemental Figure 2.**
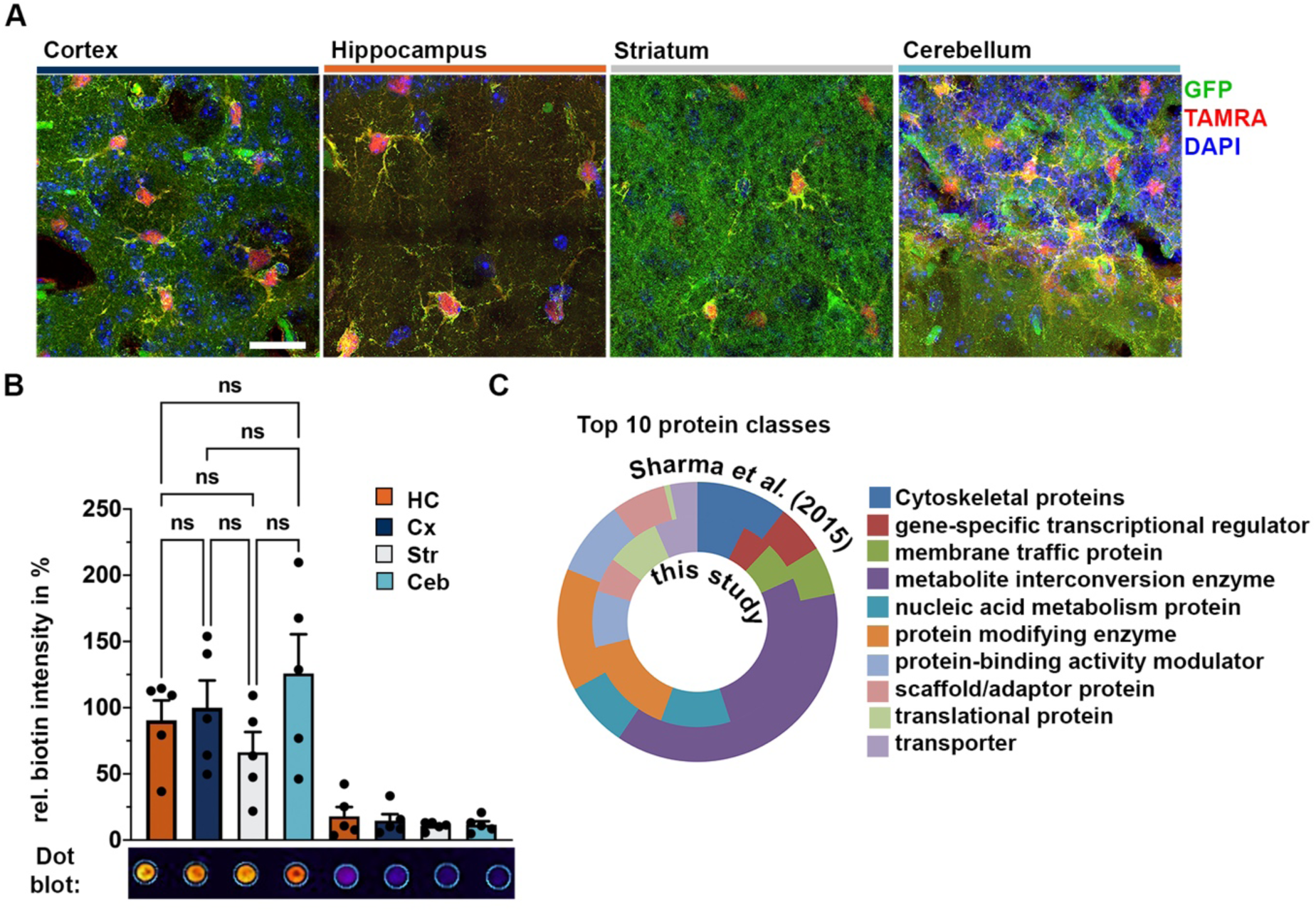
Validation of astrocyte-specific labeling. **(A)** Brain tissue slices of Cre+ MetRS^L274G^ mice were prepared and processes as described in Figure 1. TAMRA signal is evident and specific for astrocytes that also express the GFP reporter in all four brain regions used for proteomic analysis. Cells display typical astrocytic morphology (scale=20µm). **(B)** Biotin intensity of samples from Figure 2D were quantified by dot blot analysis. (n=5/group, tr=2; One-Way ANOVA (F(7)=9.498, p<0.0001); multiple comparison; n.s.: not significant; only comparison of Cre+ samples are shown). **(C)** PANTHER was used to compare glia proteins extracted from Sharma et al. with the cortex dataset of 3-fold enriched proteins (Sharma et al., 2015). Similar Top10 protein classes (by number of proteins) were identified in both datasets.

**Supplemental Figure 3.**
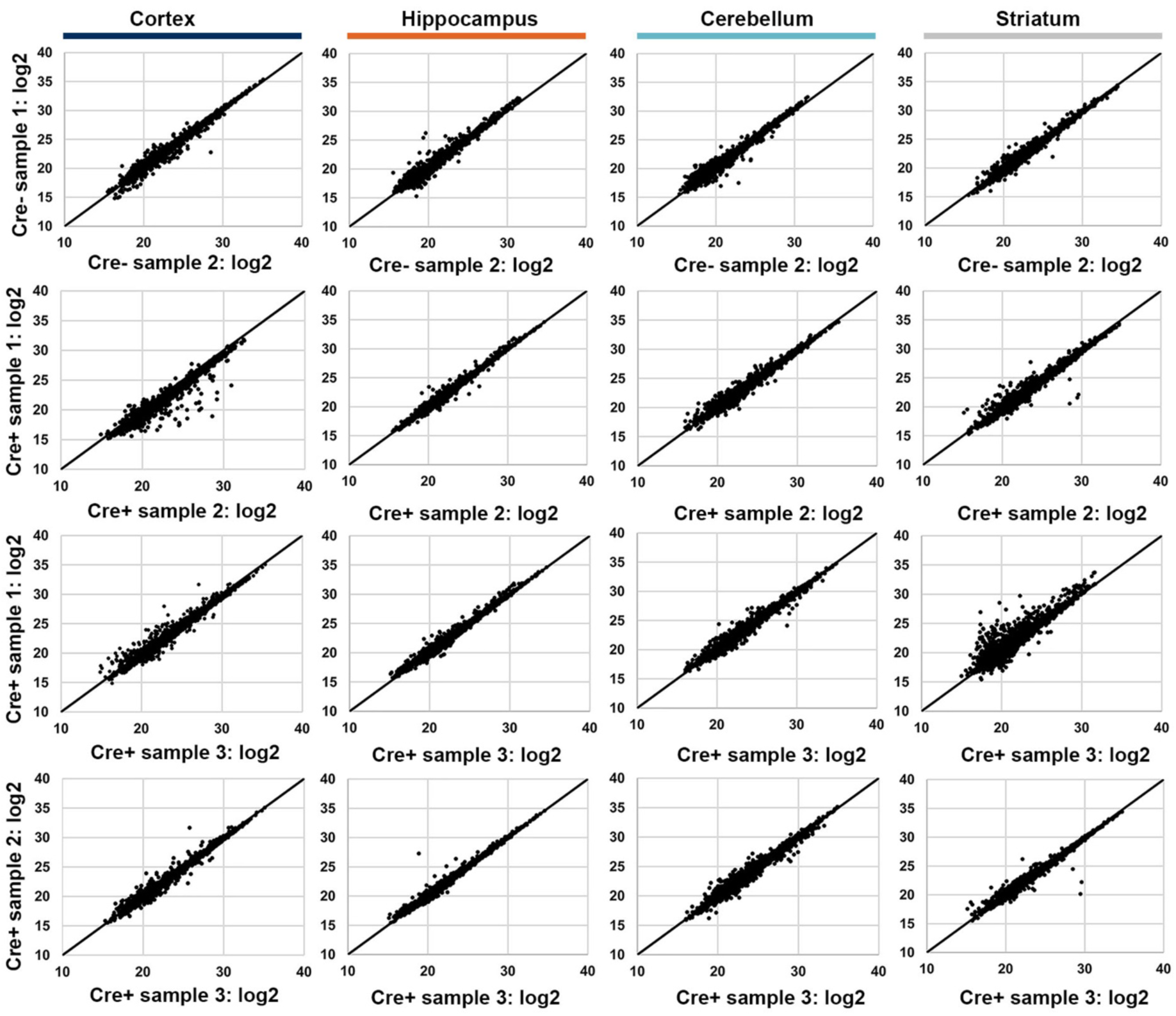
Validation of sample quality. Mass spectrometry data of each biological replicate of either Cre+ or Cre- MetRS^L274G^ derived sample was compared pairwise with each other to control for sample quality and comparability. High correlation is evident among log2 intensities of all samples in all four brain regions tested.

**Supplemental Figure 4.**
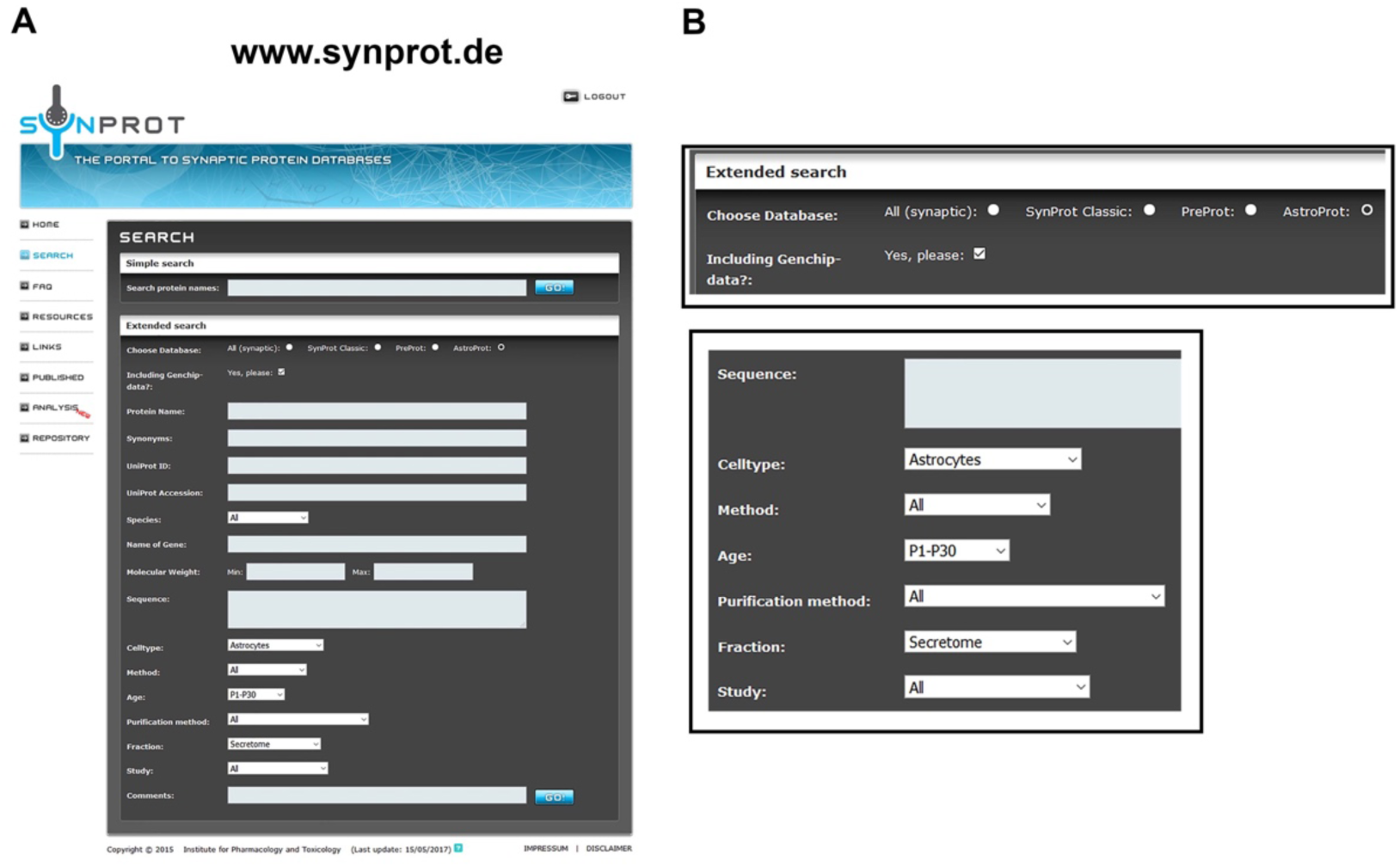
Screenshot of AstroProt database. (A) AstroProt database user interface (B) Astroprot database allows to search for entries according to different parameters such as method of preparation, cell compartment or age of rodents used. Single candidate searches can be done in AstroProt as well as in SynProt or PreProt that focusses on synaptic proteins.

**Supplemental Figure 5.**
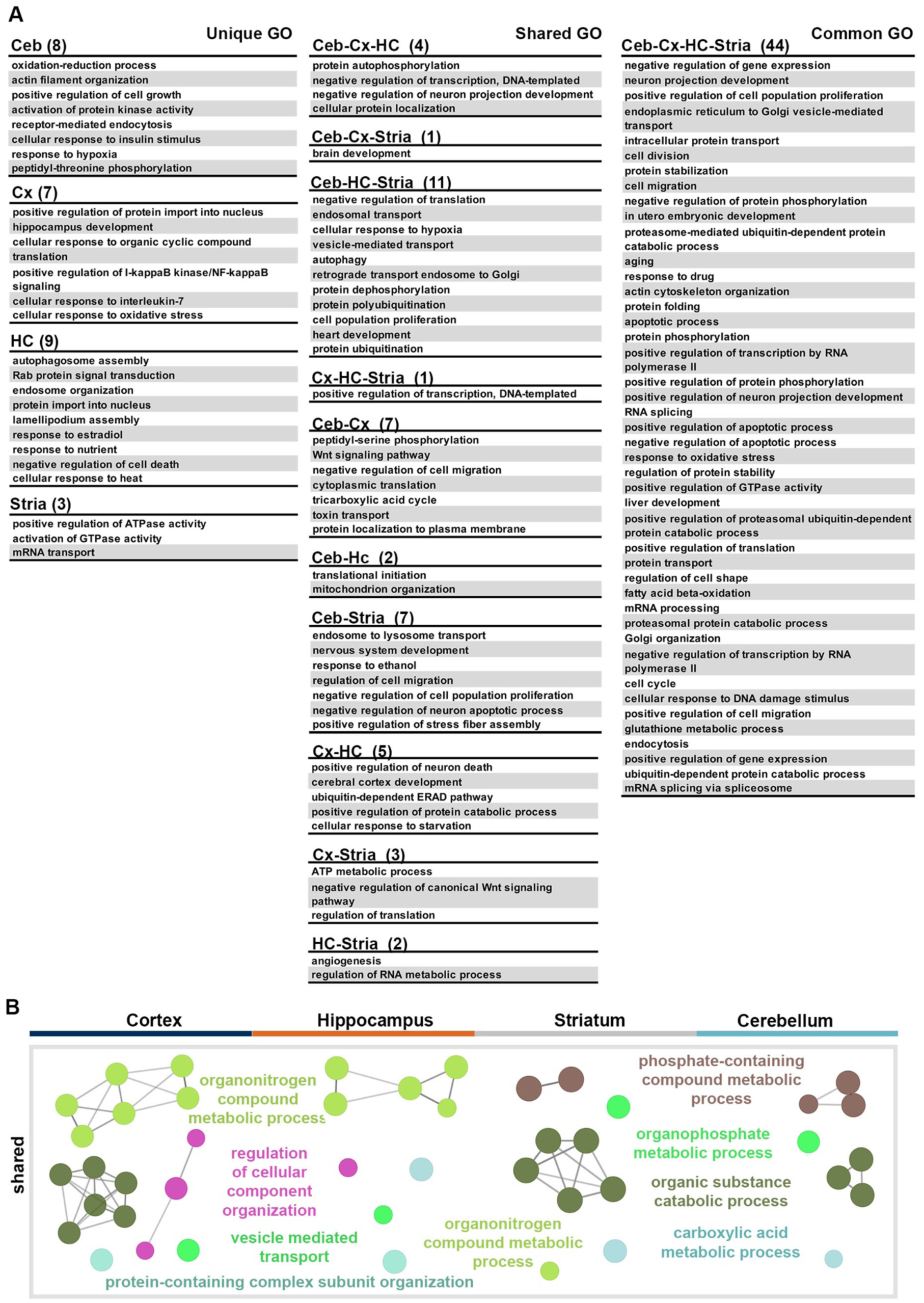
Cluego analysis: shared biological processes. (A) Lists of significant GO terms that were plotted as Venn diagram in Figure 4. GO terms are either unique for one region, common for all regions or shared by at least two brain regions. (B) Datasets of enriched proteins (≥3-fold change) for each region were analyzed by Cytoscape/ClueGO (GO biological process, only experimental evidence; p≤0.001; the node size denotes the significance). Shown here are processes that are shared by two or three brain regions: cortex (Cx), hippocampus (HC), striatum (Str) or cerebellum (Ceb).

**Supplemental Figure 6.**
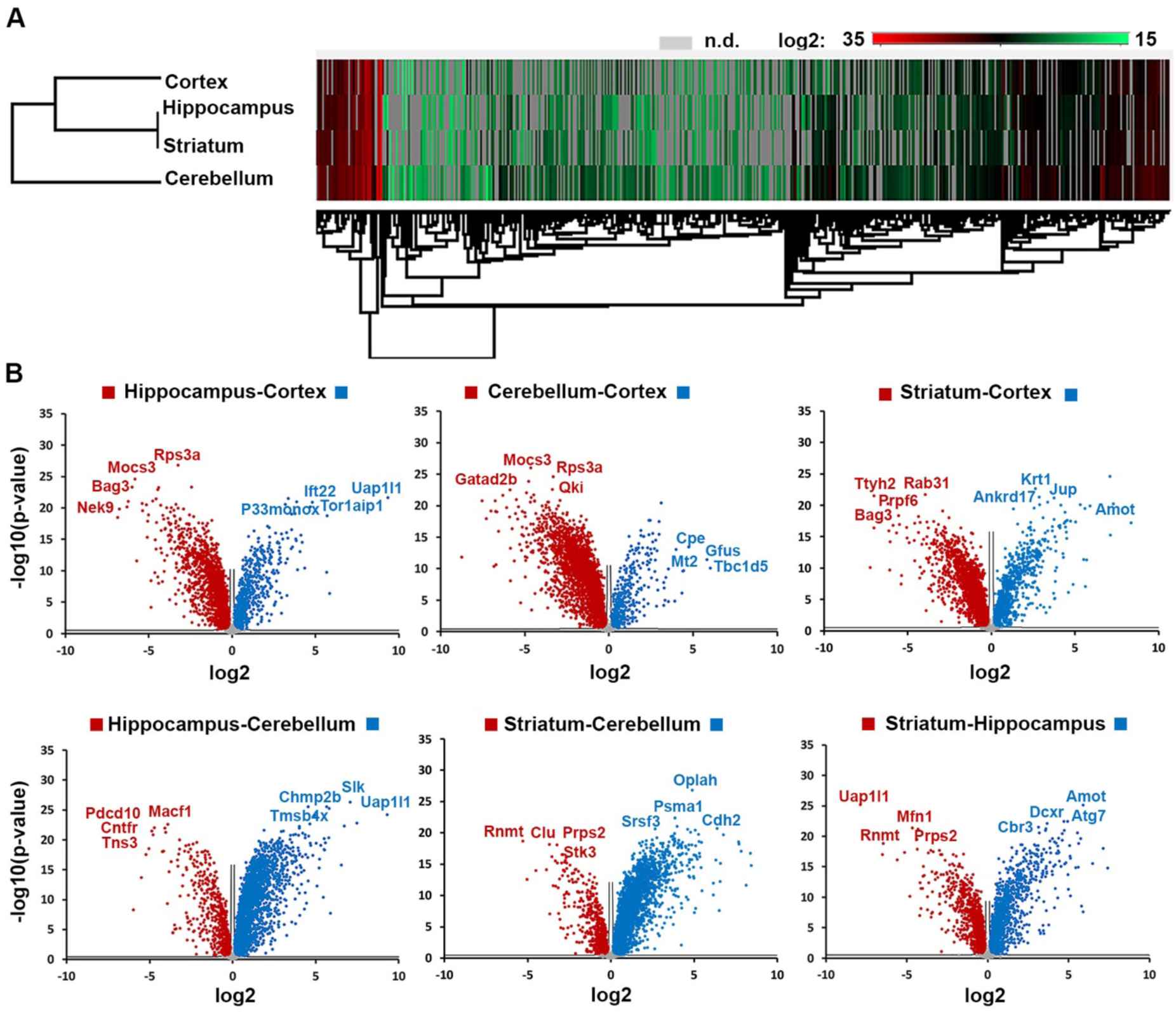
Cluster analysis and volcano plots. (A) Cluster analysis was performed with datasets from all four brain areas analyzed. For each protein, color-coded log2 intensities are displayed. Depicted here is the complete cluster analysis whereas only the top 47 proteins are shown in Figure 5A. Grey fields correspond to proteins either not detected in this region or not 3-fold enriched over Cre- MetRS^L274G^ controls. (B) Pairwise comparison of p-values plotted against high or low log2 intensity differences among pairs of datasets in volcano plots. Examples of identified candidates are displayed for all four brain regions tested showing strong intensity differences (corresponding gene name used).

**Supplemental Figure 7.**
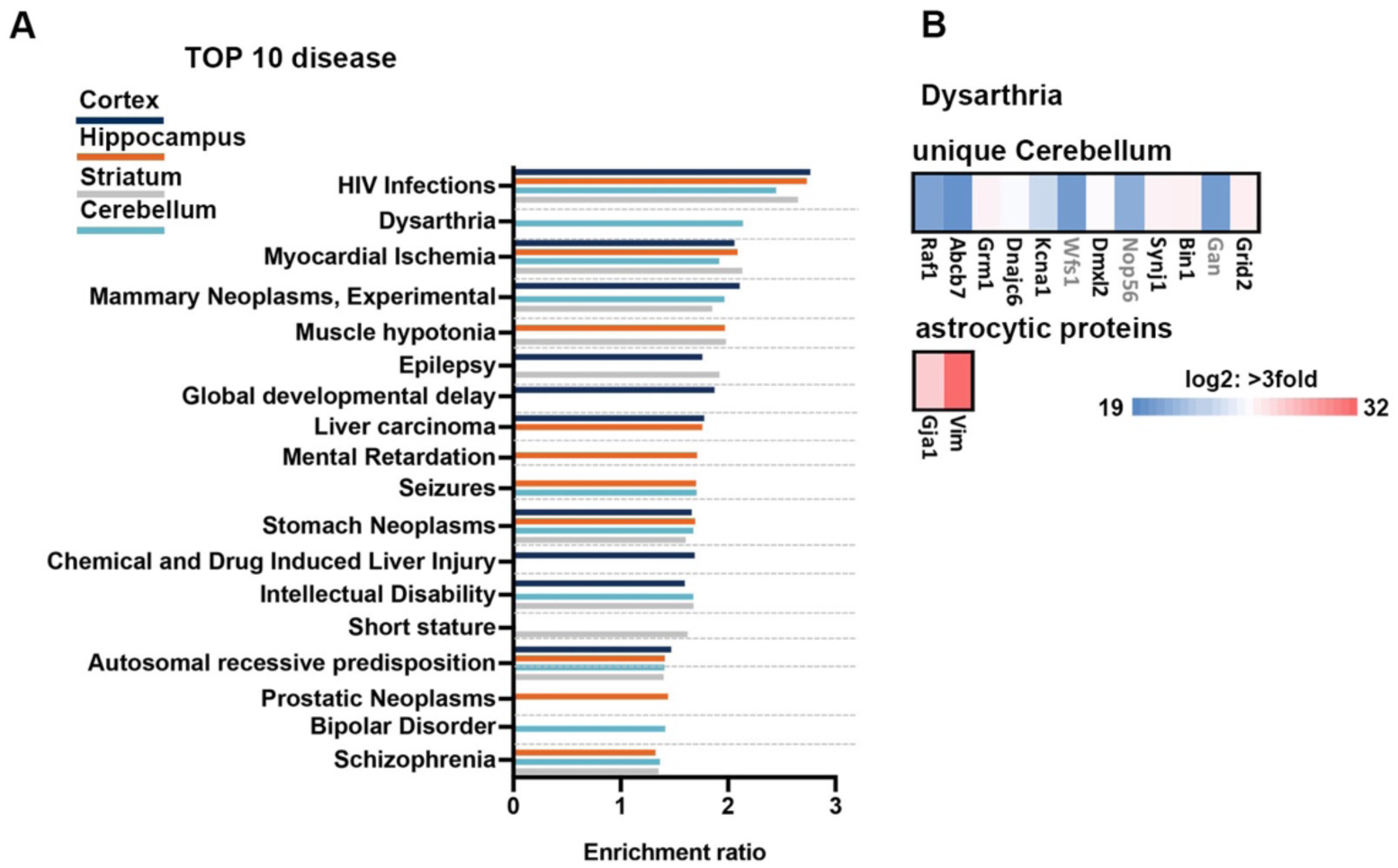
Gene enrichment analysis with disease association. (A) Gene enrichment analysis focused on diseases. Over-representation analysis was performed: DisGeNET and WebGestalt (WEB-based GEne SeT AnaLysis Toolkit). The weighted set cover algorithm summarizes each significant term into 10 categories. (B) All 89 gene names annotated to “Dysarthria” were compared with proteins uniquely identified in the cerebellum or with the list of prominent astrocytic candidate proteins from Figure 5B and plotted with their respective gene name and with color-coded log2 intensities. Gene names in black refer to reviewed protein entries in UniProt.

**Supplemental figure 8.**
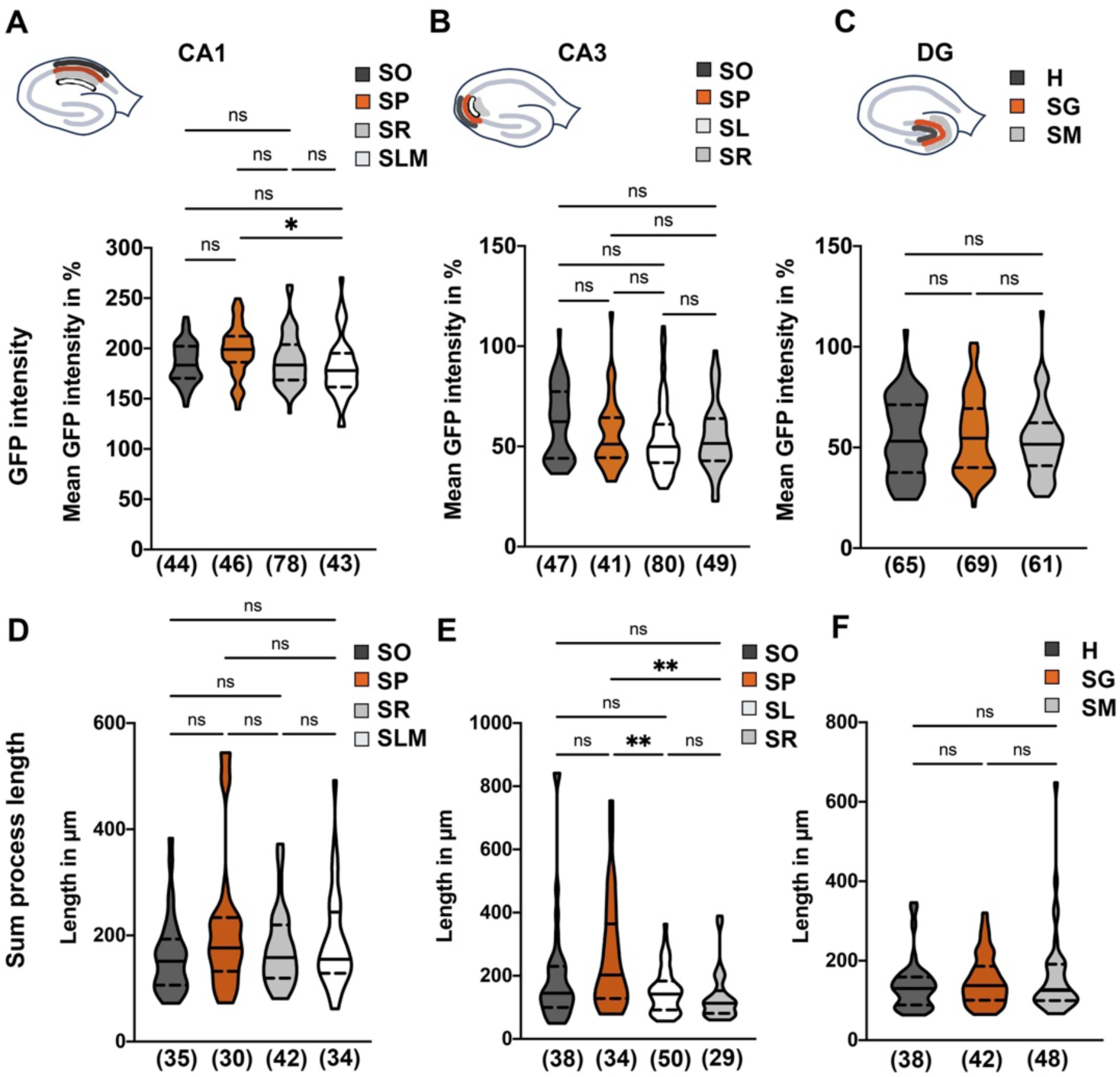
Supplemental morphological analysis of hippocampal astrocytes (A-C) Mean GFP intensity of astrocytes from sublayers of CA1 (A), CA3 (B), DG (C) was analyzed and set relative to mean GFP of all hippocampal astrocytes. (lines indicate median and quartiles; SO: stratum oriens; SP: stratum pyramidale; SR: stratum radiatum; SLM: stratum lacumosum moleculare; SL: stratum lacumosum; SM: stratum moleculare; SG: stratum granulosum; H: hilus). **(D-F)** Sum of process length is plotted for either CA1 (D), CA3 (E) or DG (F) sublayers. Astrocytes of CA3 Stratum pyramidale show a higher overall process length (E) whereas DG astrocytes process length is uniform (F). **(A-F)**: ONE-way ANOVA; A: F(3)=3.446, p=0.0176, B: F(3)=2.381, p=0.0706; C: F(E: F(2)=0.5215, p=0.5945; D: F(3)=2.111, p=0.1017; E: F(3)=6.208, p=0.0005; F: F(2)=0.7276, p=0.4853, multiple comparison (*: p<0.05; **: p<0.01) Numbers in brackets indicate number of cells used for analysis.

## Notes

### Competing Interest Statement

The authors have declared no competing interest.

